# e3SIM: epidemiological-ecological-evolutionary simulation framework for genomic epidemiology

**DOI:** 10.1101/2024.06.29.601123

**Authors:** Peiyu Xu, Shenni Liang, Andrew Hahn, Vivian Zhao, Wai Tung ‘Jack’ Lo, Benjamin C. Haller, Benjamin Sobkowiak, Melanie H. Chitwood, Caroline Colijn, Ted Cohen, Kyu Y. Rhee, Philipp W. Messer, Martin T. Wells, Andrew G. Clark, Jaehee Kim

**Author notes:** Equal contributions (Peiyu Xu, Shenni Liang, Andrew G. Clark, Jaehee Kim).

## Abstract

Infectious disease dynamics are driven by the complex interplay of epidemiological, ecological, and evolutionary processes. Accurately modeling these interactions is crucial for understanding pathogen spread and informing public health strategies. However, existing simulators often fail to capture the dynamic interplay between these processes, resulting in oversimplified models that do not fully reflect real-world complexities in which the pathogen’s genetic evolution dynamically influences disease transmission. We introduce the epidemiological-ecological-evolutionary simulator (e3SIM), an open-source framework that concurrently models the transmission dynamics and molecular evolution of pathogens within a host population while integrating environmental factors. Using an agent-based, discrete-generation, forward-in-time approach, e3SIM incorporates compartmental models, host-population contact networks, and quantitative-trait models for pathogens. This integration allows for realistic simulations of disease spread and pathogen evolution. Key features include a modular and scalable design, flexibility in modeling various epidemiological and population-genetic complexities, incorporation of time-varying environmental factors, and a user-friendly graphical interface. We demonstrate e3SIM’s capabilities through simulations of realistic outbreak scenarios with SARS-CoV-2 and *Mycobacterium tuberculosis*, illustrating its flexibility for studying the genomic epidemiology of diverse pathogen types.

## 1 Introduction

Infectious disease dynamics involve complex interactions among epidemiological, ecological, and evolutionary (epi-eco-evo) processes [1–3]. For instance, mutations to a pathogen’s genome can affect its transmissibility [4, 5] or drug resistance [6, 7], impacting epidemic progression and the efficacy of intervention strategies. To accurately represent these complexities, an integrated computational approach is essential for modeling and analyzing genetic, ecological, and epidemiological data [8, 9]. Traditional analytical methods are often insufficient to capture these complexities [10, 11], necessitating the development of more advanced simulation tools.

Simulations have become indispensable for investigating analytically intractable models [12, 13] and validating statistical inference methodologies [14, 15]. In genomic epidemiology, they provide a controlled platform to understand the complex relationships between disease transmission, pathogen evolution [16, 17], and environmental factors [18], as well as for evaluating intervention strategies [19–21] and predicting future outbreaks [22]. Simulations also facilitate the development and testing of hypotheses [23], algorithms [24], and models [25], particularly when real-world data with known ground truth are unavailable, as is often the case in epidemiological, ecological, and evolutionary studies [26]. High-quality synthetic datasets are especially crucial for deep learning models, which are increasingly used in genomic epidemiology [27–30], as these models require extensive, accurate data with known ground truth to effectively learn and predict epidemic dynamics [31, 32].

Existing simulators [33–41] for genomic epidemiology have provided valuable insights into epidemic dynamics and pathogen evolution. However, they fail to address the complex interplay between disease dynamics and pathogen genetic evolution. These simulators typically follow a two-step process: first, they simulate the entire epidemic as a transmission tree; then, they superimpose the mutational process onto the resulting genealogical structure to generate synthetic pathogen sequence data (Figure 1A). While computationally efficient, this approach inherently assumes the independence of evolutionary processes and epidemic dynamics, ignoring their dynamic coupling (Figure 1B)—a crucial factor in accurate epidemiological modeling. This is especially important if one wants to develop methods to infer changes in pathogen transmission from the topology of genealogy.

**Figure 1.**
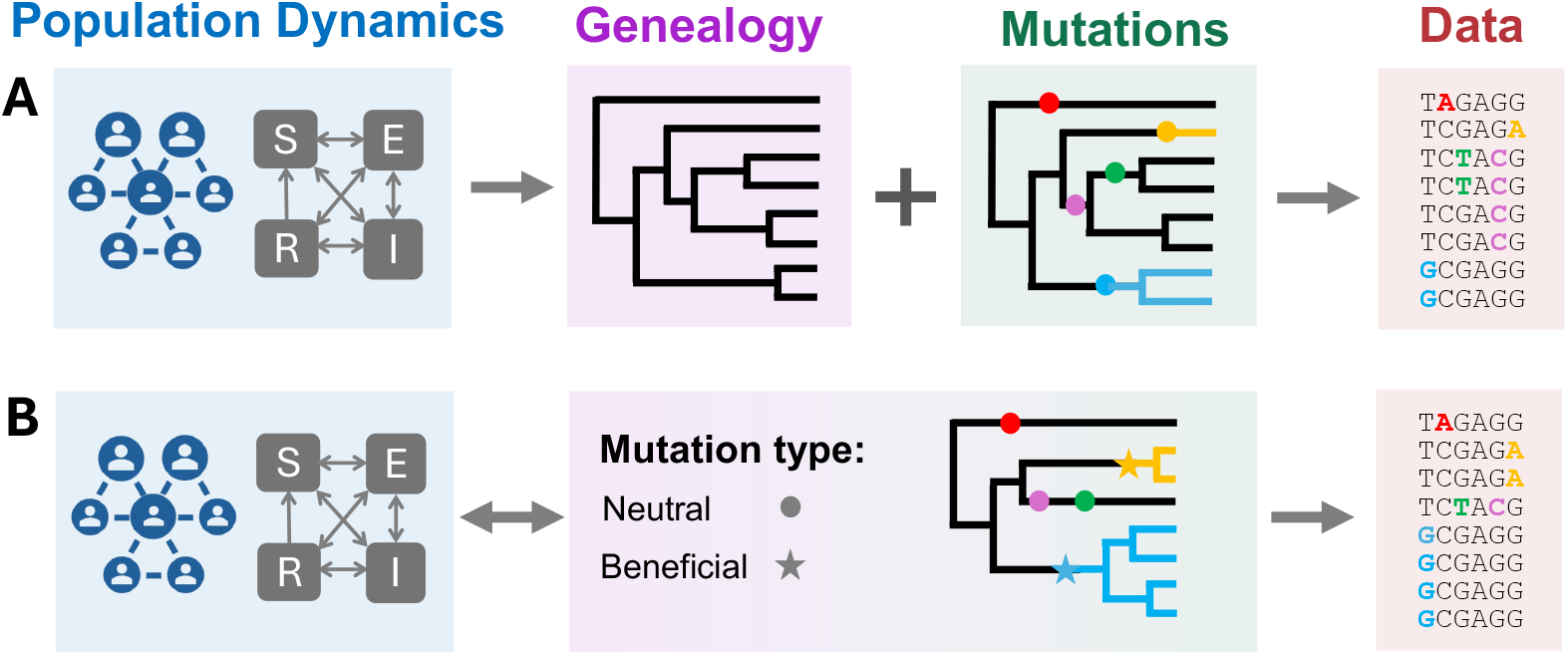
Schematic overview of the coupling of epidemiological, ecological, and evolutionary (epi-eco-evo) processes in genomic epidemiology simulators. (A) Conventional frameworks assume that genealogy generation is independent from the evolutionary process. A genealogy is first generated from epidemiological and ecological dynamics, such as the host contact network and compartmental model. Mutations are then superimposed onto the genealogy, implicitly assuming neutrality and independence between epi-eco-evo processes. (B) e3SIM integrates epi-eco-evo processes. The evolutionary process directly influences tree generation by altering the pathogen’s genetic traits, such as transmissibility or drug resistance, thereby affecting epidemic dynamics and the sampled sequencing data. This integrated approach provides a more realistic modeling framework that captures the dynamic interplay between pathogen evolution and epidemiological spread.

We introduce the epidemiological-ecological-evolutionary simulator e3SIM, an open-source software package designed to simultaneously simulate the transmission dynamics and molecular evolution of pathogens within a host population while incorporating environmental factors. Using an agent-based, discrete, forward-in-time approach with SLiM [42] as its backend, e3SIM realistically models the complex interplay between epi-eco-evo processes (Figure 1B). Integrating a SEIRS compartmental model with a host population contact network, e3SIM applies principles from epidemiology and population genetics to accurately reflect real-world scenarios. While existing software provide some insights into the connection between pathogen evolution and epidemiological traits [35, 37, 40, 41], e3SIM stands out for its comprehensive integration of epi-eco-evo dynamics, emphasizing the influence of genetic factors on these interconnected processes. Its modular design ensures flexibility and scalability, supporting various compartmental and population-genetics models, sampling schemes, and complexities such as latent infections and within-host evolution. Equipped with a graphical user interface (GUI), e3SIM provides a user-friendly and comprehensive simulation framework for genomic epidemiology.

## 2 New Approaches

e3SIM comprises three components: (1) pre-simulation modules for configuring the simulation (Section 2.1); (2) a main module for executing the simulation (Section 2.2); and (3) post-simulation modules for analyzing and visualizing the simulated results (Section 2.3). The workflow begins with the sequential execution of the pre-simulation modules, as illustrated in Figure 2, which configure the main simulation module. To streamline this process, a GUI (Section 2.4) is available for users who prefer visual interaction over command-line tools. The main module then employs SLiM [42] as the backend to perform the epi-eco-evo simulation, with its components outlined in Figure 3A–C. Finally, the post-simulation modules produce visualizations and metadata files of the simulation results, as shown in Figure 3D.

**Figure 2.**
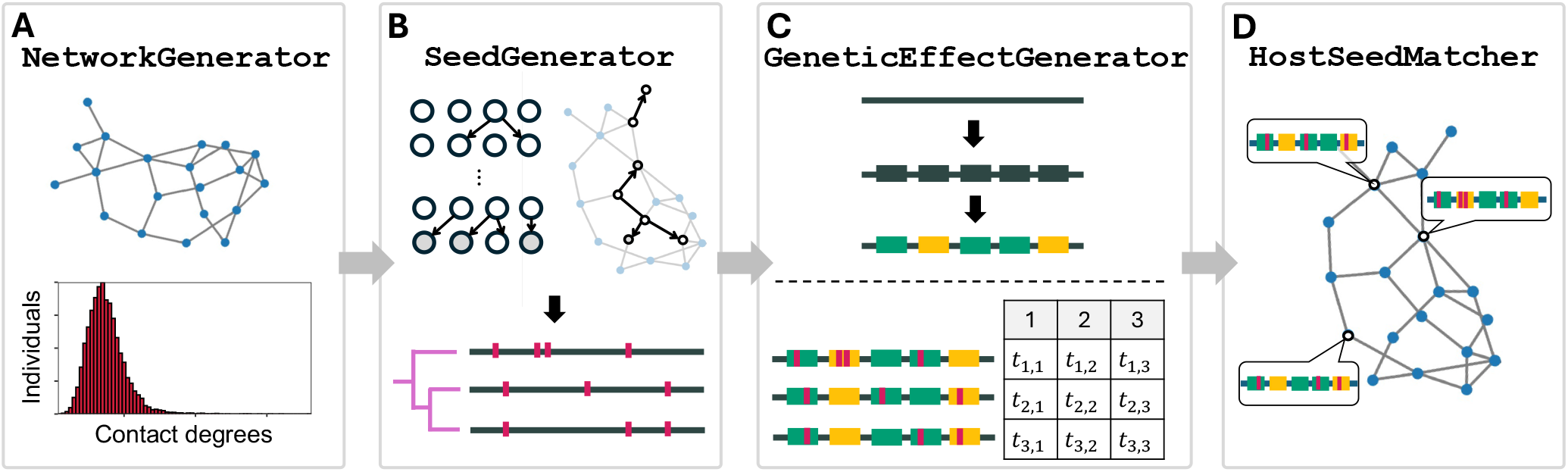
Pre-simulaton modules. A summary of the input, output, and functions of each module is provided in Table S1. (A) `NetworkGenerator` (Section 2.1.1): Constructs a host contact network, where each host is represented by a node and contacts between hosts are represented by edges. The histogram shows the distribution of contact degrees among individuals in the network. Users also have the option to input a custom contact network. (B) `SeedGenerator` (Section 2.1.2): Generates initial pathogen genomes (“seeds”) and their phylogeny using either a Wright–Fisher model (top left) or a network-based epidemiological model (top right). Pathogen genomes are depicted as horizontal lines, with vertical lines indicating mutations. Alternatively, users can provide seeds with a `VCF` file. (C) `GeneticEffectGenerator` (Section 2.1.3): Defines functional genomic regions and assigns effect sizes for traits such as transmissibility and drug resistance to mutations within these regions. These effects can be positive (green), negative (yellow), or zero for each trait. Trait values for each seed are calculated as the sum of the effects of its mutations. In the table, *t*_*i, j*_ denotes the *j*-th trait of the *i*-th seed pathogen. (D) `HostSeedMatcher` (Section 2.1.4): Assigns seed pathogens to hosts within the contact network, initiating the simulation of pathogen spread.

**Figure 3.**
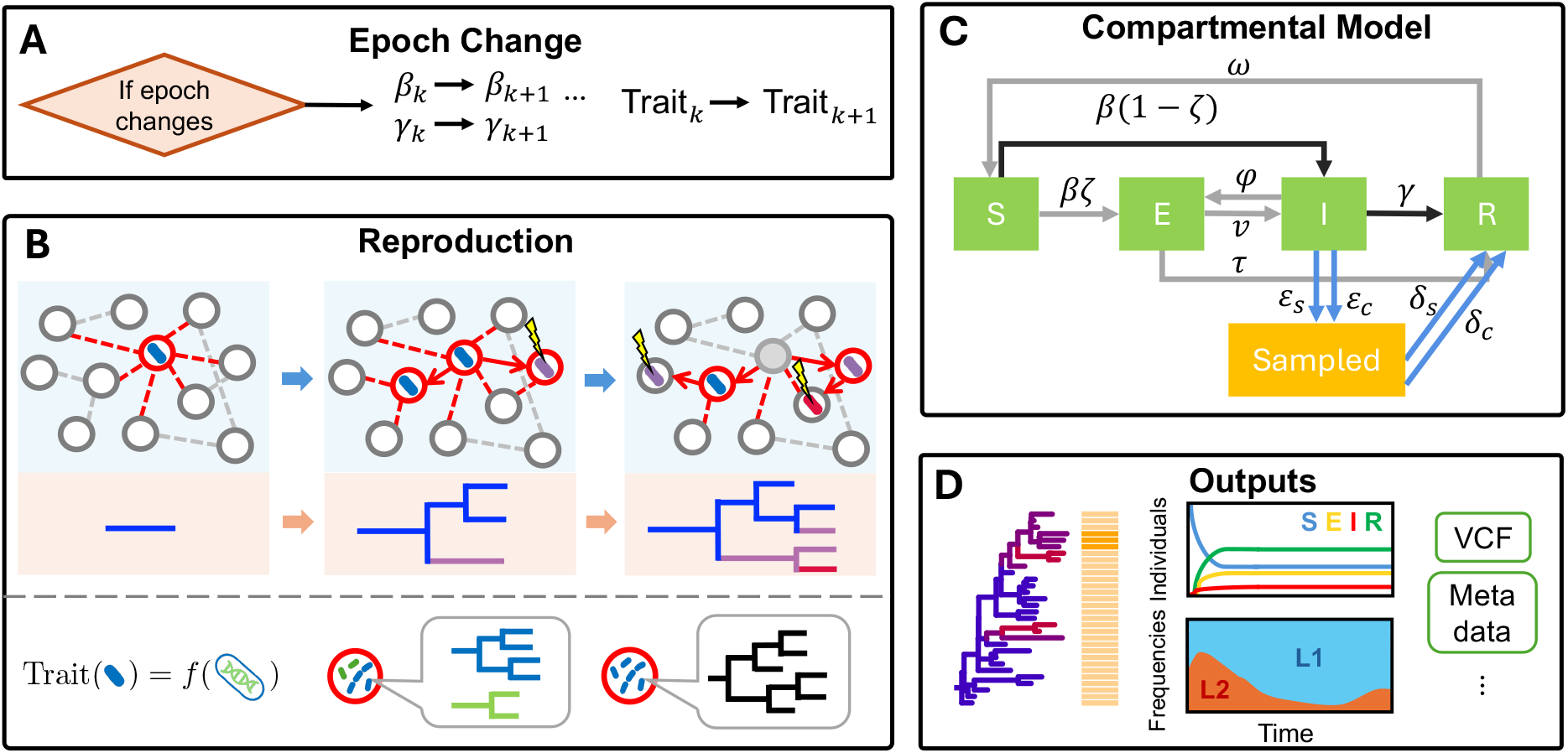
Main simulation scheme for e3SIM. Details of the modules in the simulation and an overview of the relevant configurations are listed in Table S2. (A) **Epoch change** (Section 2.2.1). Epoch changes occur as the first event in a simulation tick if that tick is scheduled for an epoch-changing event. When transitioning from the *k*-th epoch to the (*k* + 1)-th epoch, epidemiological parameters shown in panel C, such as the transmission probability (*β*_*k*_) and recovery probability (*γ*_*k*_), are updated to their respective new values. Additionally, the genetic architecture for each trait, such as transmissibility or drug resistance, is updated according to the specifications for the new epoch. This mechanism allows the simulation to incorporate time-dependent changes in environmental and epidemiological conditions. (B) **Reproduction** (Section 2.2.3). Reproduction is the second event in a simulation tick. The top row depicts the transmission process over two successive ticks. Transmission occurs over a contact network with a probability determined by both the base transmission probability and the transmissibility trait values, represented by the branch color of the genealogy (blue: low, red: high). The trait values are calculated based on the mutation profiles of the pathogen genome (panel (i)) using the genetic architecture (Section 2.1.3). Mutations, indicated by the yellow lightning symbol in the top row, are introduced concurrently with reproduction. Super-infection (panel (ii)), which involves a host being infected by multiple strains simultaneously, and within-host replication (panel (iii)), which refers to the replication of the pathogen within a single host, can also occur if specified by the user. (C) **Compartmental model and sampling** (Sections 2.2.4 and 2.2.5). Compartmental transition decisions are made for all hosts and pathogens based on the probabilities of transitioning to other states. The diagram shows all permissible transitions, with black lines indicating mandatory transitions and light grey lines indicating user-specified optional transitions (Section 2.2.4). Blue lines represent sampling events as described in Section 2.2.5, with probabilities *ε*_*s*_ for sequential sampling and *ε*_*c*_ for concerted sampling, both leading to recovery with probabilities *δ*_*s*_ and *δ*_*c*_, respectively. The state changes of hosts are processed following the completion of all reproduction events. All symbols for the epidemiological parameters are listed in Table S3. (D) **Outputs** (Section 2.3). After the simulation, several output files are generated, including visualizations of the genealogy (left), time-trajectories of host compartment sizes (middle top), lineage proportion trajectory (middle bottom), `VCF` files of sampled pathogens, and metadata files (right). When a genetic architecture is used, branches are color-coded according to user-selected trait values, and the visualization includes a heatmap displaying the user-selected trait values. These outputs provide comprehensive insights into the simulation results, enabling detailed analysis of epidemiological and evolutionary dynamics.

### 2.1 Pre-simulation modules: input file generation

Before initiating the pre-simulation modules, users must provide a pathogen reference genome in `FASTA` format, which serves as the basis for defining the genetic architecture and annotating mutations. The four pre-simulation modules (Figure 2) should then be run sequentially as prerequisites for the main module. Table S1 summarizes the input and output files for each pre-simulation module. Users can either customize and provide these files directly, or use versions that are randomly generated by e3SIM.

#### 2.1.1 Host population contact network

The host contact network is modeled as an undirected, unweighted graph, with nodes representing hosts and edges indicating contacts between them (Figure 2A, top). In e3SIM, transmissions occur only between directly connected hosts, making the contact network crucial for modeling outbreak dynamics [43]. The network structure remains static throughout the simulation. The host population contact network must be provided as a tab-delimited adjacency list. Users can supply a custom network in this format or use the `NetworkGenerator` module to generate a random contact network for a specified population size using one of several supported random network models (Figure 2A). e3SIM uses NetworkX [44] for backend random network generation, supporting Erdős–Rényi [45], Barabási–Albert [46], and random-partition [47] networks. Based on the selected random network model and parameters, `NetworkGenerator` generates a contact network and stores it in the user-defined working directory. The Erdős–Rényi network models a well-mixed host population with a normal distribution of contact degrees. The Barabási–Albert network produces a right-skewed degree distribution, indicating high heterogeneity in contact structure, suitable for modeling epidemics with “superspreaders” [33]. The random-partition network generates networks with clustering structures, suitable for simulating contact patterns in structured populations, such as regions with both rural and urban areas.

Because contact-network features strongly influence outbreak probability and outcomes [48], the specific attributes of the generated network are critical. The pre-simulation module’s GUI therefore provides a visualization of the degree distribution of the selected contact network (Figure 2A, bottom), enabling users to adjust parameters affecting network properties, such as node degree distribution and network connectivity, to ensure that the network accurately represents the desired epidemiological scenario. Although the GUI visualizes only the degree distribution, additional network characteristics, including degree-degree assortativity and concurrency, are implicitly modeled based on the underlying random graph model employed. Users also have the flexibility to customize and integrate any specific network structure, such as age-assorted networks, into the simulation via the user-input feature.

#### 2.1.2 Generation of seeding pathogen sequences

To initiate an epidemic simulation, the simulator requires “seeds”, which are the pathogen genomes infecting hosts at the start of the simulation. These genomes are represented by `VCF` files containing their mutation profiles. Users can either supply their own seed files or use the `SeedGenerator` module to randomly generate the desired number of seeds. e3SIM uses SLiM 4.1 [42] for random generation, offering two modes: Wright–Fisher mode and network-based epidemiological mode. In Wright–Fisher mode (Figure 2B, top left panel), `SeedGenerator` performs a Wright–Fisher simulation [49, 50], using a user-specified mutation rate and effective population size, starting from the reference genome and running for a specified number of generations. In network-based epidemiological mode (Figure 2B, top right panel), `SeedGenerator` simulates the initial outbreak on the contact network generated by `NetworkGenerator`, using initial pathogen genomes identical to the reference genome. In both modes, a specified number of seeding sequences for the main simulation module are randomly sampled from the final generation of simulated pathogen genomes (Figure 2B, bottom right) and stored in `VCF` files. Their genealogy is also recorded (Figure 2B, bottom left) and saved in `NWK` format.

#### 2.1.3 Genetic architecture

One of the key features of e3SIM is its capability to integrate epi-eco-evo dynamics. This integration is achieved through explicit modeling of pathogen genetic variation, which underlies key epidemiological traits such as transmissibility and drug resistance. This approach enables dynamic feedback between the evolution of these genetic traits and the epidemiological dynamics within ecological contexts. If the epi-eco-evo coupling is not required, users can opt for a trait-less model, which simulates the spread of neutrally evolving pathogens within a contact network.

The genetic architecture of pathogen traits, such as transmissibility, can be user-defined or randomly generated by the `GeneticEffectGenerator` module. Using a `GFF` (general feature format) file with reference genome annotations, typically from NCBI [51], this module selects genomic regions for each trait and assigns effect sizes uniformly sampled from a user-defined range in random generation mode. Genetic architectures for multiple traits, such as transmissibility and drug resistance, can be specified. In a dynamically changing environment, the genetic architecture can be modeled using epoch-specific frameworks by selecting the appropriate genetic architecture for each epoch. Pleiotropy, where a single gene influences multiple traits, is common in pathogen drug resistance [52] and can be incorporated by assigning different effect sizes to the same locus for multiple traits. `GeneticEffectGenerator` calculates trait values for all seeds based on the defined genetic architecture and stores them in a `CSV` file in the working directory, as shown in Figure 2C.

The overall trait value of a pathogen genome is determined by the cumulative effect of all mutations within its causal genomic elements. These elements are specific regions crucial for the pathogen’s functionality and pathogenesis, where mutations significantly impact pathogen traits. Causal genomic elements are defined by their positions within the genome, ranging from a single base pair to an entire gene, along with their associated effect sizes. For a pathogen genome *i* with *n* mutations in causal genomic elements, the trait value is given by the sum of the effects of all mutations [53, 54]:

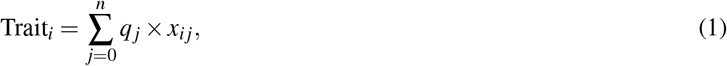

where *q* _*j*_ represents the effect size of mutation *j*, and *x*_*ij*_ is a binary indicator variable that equals 1 if mutation *j* is present in pathogen genome *i* and 0 otherwise. When using the random generation mode in `GeneticEffectGenerator`, e3SIM also includes an optional heuristic approach for normalizing the expected average trait value to a user-defined value by the end of the simulation (Section 5.1).

We define *transmissibility* as the trait influencing the probability of pathogen transmission per contact and *drug resistance* as the trait affecting the probability of pathogen survival during host treatment. For a pathogen *i*, the per-contact transmission probability and the per-pathogen survival (i.e., treatment failure) probability within a tick are computed as follows:

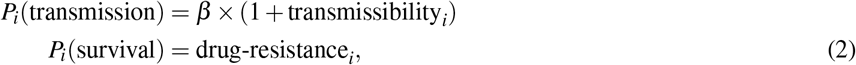

where *β* is the base per-contact transmission probability (Figure 3C).

#### 2.1.4 Seed-host matching

To start a simulation in e3SIM, the initial host(s) (“patient zero”) must be seeded with pathogen genomes (Figure 2D). This process requires providing the main simulator module with a `CSV` file that specifies the mapping between seeds and hosts. This file can be either generated by the `HostSeedMatcher` module or user-provided. Note that there may be more than one initial host. `HostSeedMatcher` offers three schemes for matching seeds to hosts. The “Uniform Selection” scheme assigns each seed to a randomly selected host from the pool of all available hosts. The “Contact Ranking” scheme assigns seeds based on the host’s contact degree ranking; for example, the host with the highest number of contacts receives the first seed, the second-highest receives the second seed, and so on. Each seed can be assigned to a host with any rank. The “Percentile-Based Selection” scheme assigns seeds based on a specified percentile of the contact degree distribution, such as selecting hosts from the top 10% with the highest number of contacts. Different schemes can be applied for each seed. When using the GUI for pre-simulation configuration, users can view the degree distribution of the host contact network, select a matching method for each seed, and inspect the matching results interactively within the contact network’s degree distribution.

### 2.2 Main module: epi-eco-evo simulation

After completing all pre-simulation modules, the `OutbreakSimulator` module of e3SIM can be executed using a configuration file in `JSON` format. `OutbreakSimulator` leverages SLiM as the backend and generates a script written in Eidos, SLiM’s scripting language. This script is assembled from Eidos code blocks based on the specified configuration parameters, implementing the epi-eco-evo simulation specified by the user. Given user choices for functions such as within-host replication and super-infection, `OutbreakSimulator` includes the necessary script blocks and schedules them at specific time points during the simulation. Multiple replicates with the same configuration can also be conducted. Below, we provide a detailed description of e3SIM’s simulation procedure (Figure 3A–C). Advanced SLiM users can modify the generated Eidos script stored in the working directory to extend the dynamics of the simulation in ways not directly supported by e3SIM.

For the epidemiological model, e3SIM employs a flexible SEIRS compartmental framework [55–57]. Hosts are classified into one of four compartments: Susceptible, Exposed, Infected, or Recovered, with various state transitions (Figure 3C).

All pathogens are modeled explicitly in e3SIM as “individuals”, and all hosts are modeled as “subpopulations” in SLiM’s terminology [42]. Transitions between compartments are governed by base probabilities specified in the configuration file, modified by traits of the pathogens in the host. This agent-based approach simulates dynamics for each host rather than relying solely on population averages. Adjusting the transition probabilities allows the simulation of various compartmental models, such as SIR and SEIR, enabling flexible modeling of diverse disease dynamics and transmission scenarios. The following sections detail each transition between compartments and their integration within the epi-eco-evo simulation process in e3SIM.

#### 2.2.1 Ticks and epochs

`OutbreakSimulator` operates in discrete time, where each step, or *tick*, represents the unit of time during which actions occur for each pathogen and host. The simulation concludes when a predefined number of ticks is reached. In every tick, decisions are made for all possible reproduction and compartmental transitions. Ticks are organized into *epochs*, each comprising a sequence of consecutive ticks. Within each epoch, the base rates for compartmental transitions and the genetic architecture for each trait remain constant. These parameters can change to new values when the simulation transitions to a new epoch, as shown in Figure 3A. The epoch structure allows the simulation to incorporate time-dependent environmental changes, such as implementing new intervention strategies at specific time points after the outbreak starts.

#### 2.2.2 Evolutionary model

Mutations accumulate in all existing pathogen genomes at each time tick according to a user-specified substitution model. The transition probability matrix **P** for nucleotides is a 4 × 4 matrix where *P*_*ij*_ (*i* ≠ *j*) represents the probability of a nucleotide in state *i* transitioning to state *j* in one time tick (*i, j* ∈ {A, C, G, T}). The overall probability of a site in state *i* acquiring a mutation in each tick is given by ∑ _*j*≠*i*_ *P*_*ij*_. If no transition probability matrix is provided, users must specify a mutation rate, defaulting to the Jukes–Cantor model [58]. Mutations occur stochastically at a constant probability each tick, as defined by the mutation probability matrix, regardless of whether the pathogen reproduces. Trait values, influenced by the mutation profiles of the pathogens (Section 2.1.3), are recalculated before transmission events (Figure 3B(i)). These recalculated trait values subsequently alter the probabilities of relevant transitions, such as transmission and recovery. This mechanism captures the crucial interaction between epi-eco-evo processes, which is often overlooked in classical epidemiological simulators (Figure 1A).

#### 2.2.3 Reproduction: transmission and within-host replication

In e3SIM, pathogen reproduction occurs through two primary modes: transmission and within-host replication. Transmission occurs when a pathogen infects a new host by producing offspring within the new host, which can only happen when the hosts are in direct contact. During each tick, all effective contacts are evaluated for potential transmissions (Figure 3B). An effective contact involves a pair of hosts connected by an edge, where one host is infected (the infector) and the other is capable of being infected (the infectee). For each effective contact, a single pathogen is randomly selected from the infector’s pathogen population. The transmission success of the chosen pathogen is determined by a Bernoulli trial with the success probability defined as the product of the base transmission probability (*β*) and the pathogen’s transmissibility trait (Eq. 2). Successful trials result in pathogen transmission to the infectee. e3SIM assumes a transmission bottleneck [59–61], where only one pathogen genome from the infector is transmitted to the infectee per successful transmission.

“Super-infection” refers to a scenario in which a host can be infected by multiple sources, either within the same tick or across different ticks (Figure 3B(ii)). If disabled, a host can only be infected if it is in the susceptible state. In this case, only one successful transmission per tick is retained, with the infecting source chosen randomly from among the potential infectors. If super-infection is enabled, the host can be an infectee regardless of existing infection, as long as the number of pathogens within the host does not exceed the specified maximum capacity. All successful transmissions to this host within a single tick are retained.

Within-host replication occurs when a pathogen reproduces within a single host (Figure 3B(iii)). This process is enabled only if the within-host replication mechanism is active and the total number of pathogens within the host does not exceed the user-defined maximum capacity. Under these conditions, each pathogen has a probability of producing exactly one offspring within the same host during each simulation tick. Mutations accumulate over time independently of the reproduction process, as detailed in Section 2.2.2. Therefore, all pathogens, regardless of their reproduction status, have the same probability of mutation per tick.

#### 2.2.4 State changes: activation, recovery, and loss of immunity

State changes occur after reproduction events. Upon successful transmission to a susceptible host, the host transitions to either an infected or exposed state. The exposed state indicates a latent infection period during which pathogens are present and may replicate within the host, but are not yet transmissible. A newly infected host enters the exposed state with a probability ζ (Figure 3C) and the infected state with a probability 1 − ζ. Exposed hosts progress to the infected state with an *activation* probability *ν* and have a probability *τ* of recovery. Conversely, infected hosts can revert to the exposed state with a deactivation probability *ϕ*, modeling a return to latency.

*Recovery* in e3SIM models the transition of both exposed and infected hosts to a recovered state. When drug resistance genetic architecture is disabled, infected hosts recover with a probability *γ* per tick, independent of the sampling event; the sampling event can also trigger recovery through treatment, as detailed in Section 2.2.5. When the genetic architecture for drug resistance is enabled, the probability of recovery is influenced by the collective resistance traits of the pathogens. In this scenario, infected hosts are evaluated for recovery with a probability *γ*, during which each pathogen *i* within the host undergoes a Bernoulli trial with a success probability *P*_*i*_(survival) (Eq. 2). Pathogens are cleared from the host if they fail to survive in the current tick. When all pathogens are cleared, the host successfully recovers and transitions to the recovered state.

e3SIM posits that hosts in the recovered state are immune to reinfection, thereby modeling acquired immunity. However, these hosts can revert to the susceptible state with a probability *ω*, allowing potential reinfection due to immunity loss. Each host undergoes at most one state transition per tick, determined by a multinomial trial that considers all possible transitions from its current state, including the option to remain in the same state. For instance, for a host in the exposed state, the probability of remaining exposed in one tick is 1 − (*ν* + *τ*), as illustrated in Figure 3C.

#### 2.2.5 Sampling

`OutbreakSimulator` supports two types of sampling methods: sequential sampling and concerted sampling, following standard birth-death-sampling frameworks [62]. Sequential sampling selects individual lineages according to a Poisson process [63, 64], whereas concerted sampling selects multiple lineages simultaneously at predetermined time points [65]. Sequential sampling, also known as heterochronous sampling, occurs with a probability *ε*_*s*_ for each infected host and takes place simultaneously with the state change events in each tick (Section 2.2.4). Sampling can lead to an individual’s recovery due to treatment, modeled by a post-sampling recovery probability *δ*_*s*_ for each sampled pathogen, independent of the base recovery probability *γ* (Figure 3C) [64, 66, 67]. In this approach, sampling takes place continuously over time and independently for each pathogen. Concerted sampling, also referred to as isochronous sampling, happens at predefined user-specified time points, immediately following state transitions. This synchronous approach involves sampling across the entire population at specific time ticks. During each concerted sampling event, infected hosts are sampled with a probability *ε*_*c*_ and subsequently recover with a probability *δ*_*c*_. Key epidemiological events, including infection, activation, recovery, and sampling, are recorded in `CSV` files accessible in the working directory. Both sequential and concerted sampling can be enabled in the same simulation. In such cases, sequential sampling is executed first, followed by concerted sampling at specific user-defined ticks.

### 2.3 Post simulation: analysis and visualization of simulated data

Upon the completion of the simulation, `OutbreakSimulator` performs post-simulation analysis on the simulated data. The output from this analysis includes various types of information (Figure 3D). First, e3SIM generates a `VCF` or `FASTA` file containing the mutation profiles of all sampled sequences for various downstream analyses. Additionally, a metadata file with trait values for all sampled pathogens is provided. A phylogenetic tree based on genetic distance and a time-scaled tree representing real-time per user-defined tick can be inferred from the sampled pathogen sequences and their corresponding sampling times using standard phylogenetic inference software, such as IQ-TREE 2 [68], RAxML-NG [69], and TreeTime [70].

The simulation also generates a tree-sequence file [71–73], which is processed by `OutbreakSimulator` to produce a `NWK` tree file containing the genealogy of sampled sequences for each seed’s progeny. Additionally, the simulation logs detailed infection events throughout the process, documenting who infected whom. This infection data can be integrated with the genealogical information to represent the transmission dynamics. The resulting tree is equivalent to the transmission tree if neither super-infection nor within-host replication is invoked. e3SIM generates a visualization of this tree in `PDF` format using the `ggtree` package [74–76] in R. When the genetic architecture is enabled, branches are color-coded based on the calculated values of a user-selected trait for visualization. The visualization is further annotated with a heatmap showing the trait values chosen by the user. If a tree of the seeds is provided, e3SIM combines this tree of seeds with the tree recorded during the transmission simulation for each seed’s progeny, producing a complete genealogy for all sampled pathogen genomes.

e3SIM generates two additional plots: the lineage distribution trajectory and the compartment trajectory. The lineage distribution trajectory shows how the proportion of progeny from each initial seed evolves over time, while the compartment trajectory illustrates the proportion of hosts in each state over time. Each simulation replicate is depicted as a distinct trajectory using the same configuration, with an averaged trajectory across all replicates, providing a comprehensive overview of the simulation outcomes.

### 2.4 Graphical user interface (GUI)

The primary method of interacting with e3SIM is through the GUI, which was designed for users of all experience levels— particularly those who prefer visual interaction over command-line tools. Developed using the Tkinter framework [77], a widely-used Python GUI toolkit, it ensures a consistent user experience across macOS and Linux. The GUI is structured to guide users step-by-step through the process of configuring and initiating simulations, providing clear and intuitive interaction with the software’s features. Users interact with sections and tabs dedicated to various aspects of the simulation setup (Figure 4), including specifying general run parameters, configuring evolutionary and epidemiological models, setting up seed populations, and defining the network model.

**Figure 4.**
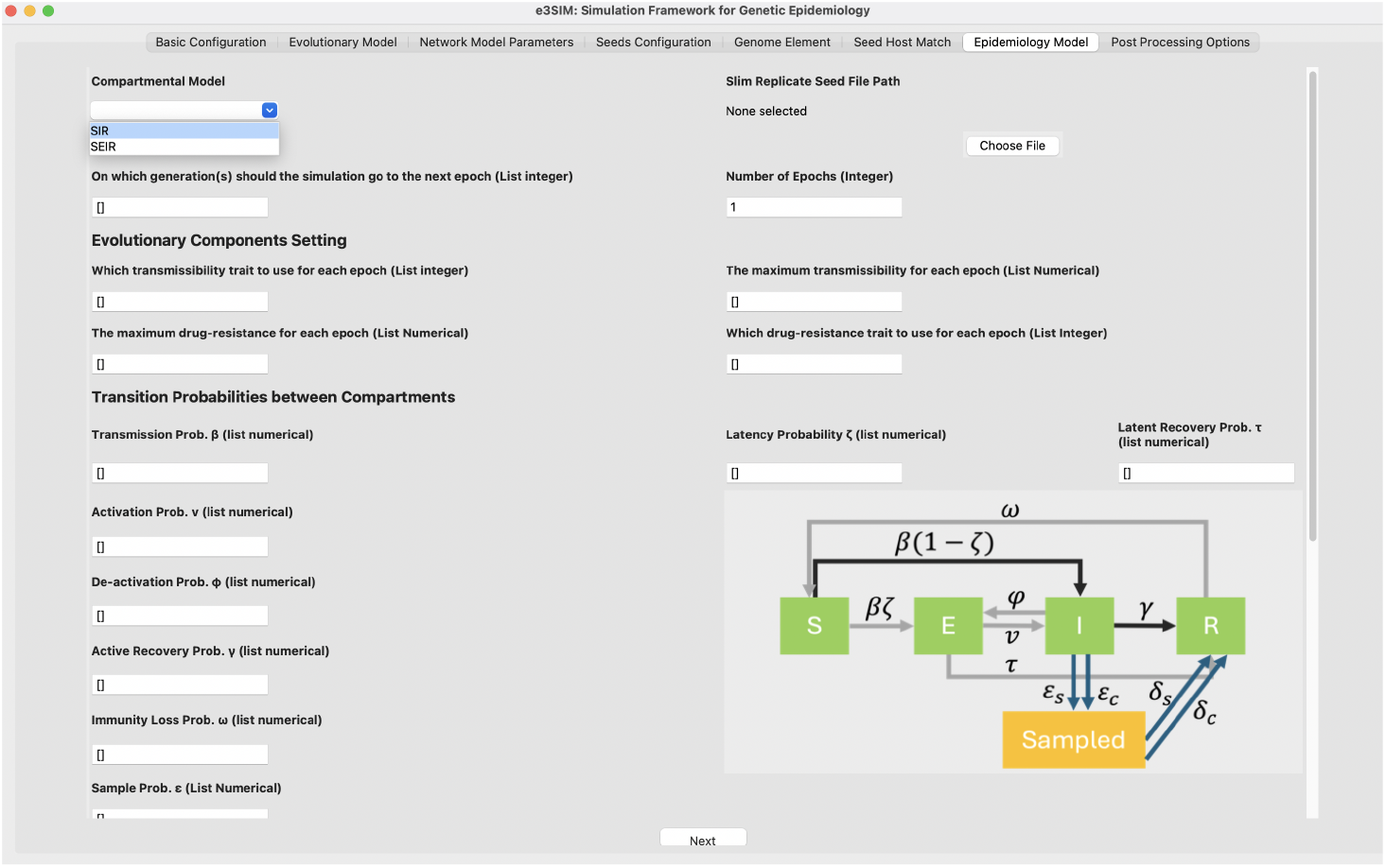
Example of the GUI for epidemiology model configuration. This tab displays the epidemiology model configuration settings, allowing users to configure the compartmental model and transition probabilities between compartments, as described in Section 2.2.4. The specified parameters are saved in a configuration file within the working directory, which can be utilized by `OutbreakSimulator` (Section 2.2).

## 3 Results

### 3.1 Simulation examples

We conducted a series of simulations demonstrating the advanced capabilities of e3SIM in capturing the complex dynamics of infectious disease spread. First, we performed simulations designed to illustrate the essential functionalities of e3SIM within the epi-eco-evo framework. Additionally, we used real epidemic datasets to model real-world outbreaks: the SARS-CoV-2 outbreak in Haslemere, UK, and *Mtb* outbreaks in Karonga, Malawi.

#### 3.1.1 Emergence of new variants and epi-eco-evo dynamics in fast-evolving pathogens

The emergence of new variants is common among fast-evolving pathogens like influenza and SARS-CoV-2 [78]. These variants may harbor mutations in functional loci that confer fitness advantages, such as increased transmissibility and resistance to treatment. We simulated a SARS-CoV-2 outbreak in a host population of 10,000 individuals using a Barabási–Albert contact network to demonstrate e3SIM’s unique capability in modeling: 1) the interplay between epidemic dynamics and adaptive genetic evolution in response to environmental changes, and 2) the interaction between disease transmission dynamics and pathogen evolution. Two distinct drug treatment strategies were applied to the entire population at different time points to show their impact on outbreak dynamics, the emergence and spread of drug-resistant strains, and the long-term evolutionary consequences of these treatment strategies. A genetic architecture was defined with one transmissibility trait and two drug resistance traits, introducing two stages of population-level treatments with different drugs, as shown in Figure 5.

**Figure 5.**
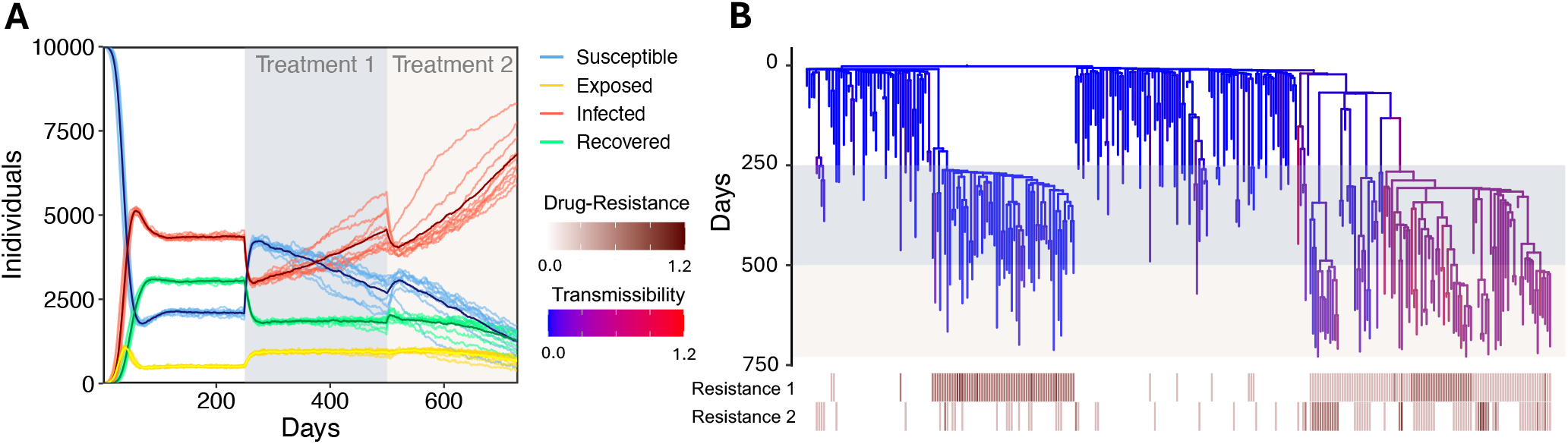
Simulated SARS-CoV-2 epi-eco-evo dynamics under two treatment stages. (A) SEIR trajectories for ten replicates of a single configuration. The bold line represents the average across all replicates. Epoch changes occur at days 250 and 500. (B) Example genealogy of sampled pathogens (*n* = 302) from one replicate produced by e3SIM. Branches are colored by the lineage’s transmissibility at each time point (blue: lowest, red: highest). Genealogies from all ten replicates are shown in Figure S1, with trees colored by resistance to the drugs shown in Figures S2 and S3. The heatmap shows the drug resistance for all the sampled individuals, with each row representing a different drug resistance trait for each sampled individual. The shaded backgrounds denote treatment stages 1 and 2 in both figures.

Before any treatment (days 0–250), the epidemic dynamics, represented by the number of individuals in different states (Susceptible, Exposed, Infected, Recovered), reached a steady state (Figure 5A), with the most pathogens exhibiting low transmissibility (Figure 5B). The first treatment, starting on day 250, initially caused a rapid decrease in the infected population and an increase in the susceptible and recovered populations, indicating the drug’s effectiveness in reducing transmission. However, the emergence of *de novo* drug-resistant mutations (Figure 5B) reduced the drug’s efficacy, leading to a gradual resurgence in the infected and exposed populations. Even after the introduction of the second treatment on day 500, the combined effects of the selective advantage of pathogens with *de novo* mutations conferring resistance to the second drug and the increased proportion of highly transmissible pathogens (Figure S4) resulted in a rise in the infected and exposed populations. These results highlight the dynamic response of the host population to sequential drug treatments, emphasizing the critical roles of drug resistance, genetic architecture, pathogen evolution, and adaptive fitness in shaping epidemic outcomes.

#### 3.1.2 Effect of superspreaders and contact network on epidemic dynamics

The presence of superspreaders—individuals who are highly socially active and thus have a higher likelihood of transmitting the pathogen—can significantly impact transmission dynamics [79]. To demonstrate e3SIM’s ability to incorporate host contact structure in outbreak modeling, we simulated three epidemic scenarios with *Mycobacterium tuberculosis* (*Mtb*) [80]: 1) random matching of seeds to hosts in a Erdős–Rényi network (Figure 6A); 2) matching high-transmissibility seeds to socially active hosts and low-transmissibility seeds to less active hosts in a Barabási–Albert network (Figure 6B); and 3) matching low-transmissibility seeds to socially active hosts and high-transmissibility seeds to less active hosts in a Barabási–Albert network (Figure 6C). Transmission dynamics were strongly influenced by the presence of socially active individuals. As shown in Figure 6D, high-transmissibility strains tended to form larger progenies, a trend amplified when these strains infected socially active hosts (Figure 6E). Conversely, low-transmissibility strains could also form large progenies when carried by socially active hosts (Figure 6F), confounding the inference of pathogen transmissibility based on cluster size [81, 82]. The size of each compartment at epidemic equilibrium and the time to reach equilibrium were also affected by the network structure and seeding strategy (Figure 6G-I).

**Figure 6.**
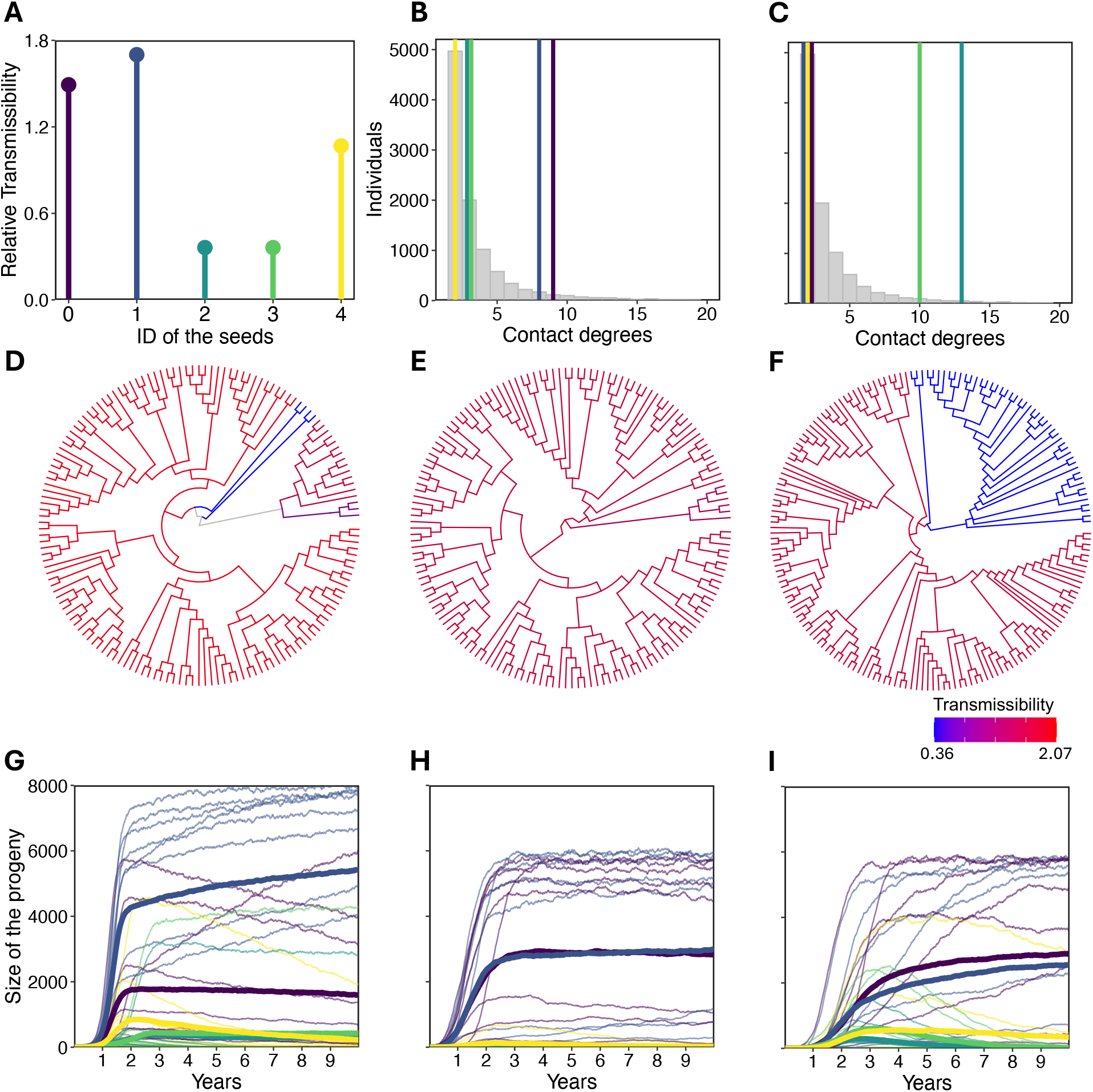
*Mtb* transmission in Erdős–Rényi network and Barabási–Albert network. (A) Relative transmissibility of the seeds (ID: 0,1,2,3,4). (B) Truncated contact degree distribution (contact degree *≤* 20) of the network used for the Barabási–Albert model simulation and the seed-host matching scheme for scenario 2. The red histogram shows the truncated contact degree distribution, and the vertical lines show contact degree of the host that is matched to each seed. The colors of the vertical lines refers to the ID of the seed as shown in Figure 6A. (C) Truncated contact degree distribution (contact degree *≤* 20) of the network used for the Barabási–Albert model simulation and the seed-host matching scheme for scenario 3. The red histogram shows the truncated contact degree distribution, and the vertical lines show contact degree of the host that is matched to each seed. The colors of the vertical lines refers to the ID of the seed as shown in Figure 6A. (D-F) One example of the simulated phylogenetic tree for sampled individuals for the three scenarios (Erdős–Rényi network with random seed matching, Barabási–Albert network with seeds matched as Figure 6B, and Barabási–Albert network with seeds matched as Figure 6C.) respectively, colored by relative transmissibility (blue: lowest; red: highest). (G-I) Trajectory of the size of the progeny for each seed for Erdős–Rényi network with random seed matching, Barabási–Albert network with seeds matched as Figure 6B, and Barabási–Albert network with seeds matched as Figure 6B. The color of each curve is matched to the ID of the seeds indicated in Figure 6A. Each curve represents one simulation replicate, with the bold curve indicating the average of the ten replicates. The SEIR trajectories corresponding to each scenario are shown in Figure S5.

#### 3.1.3 SARS-CoV-2 epidemic dynamics in Haslemere, UK: transmission of variants of concern

Between October 2023 and March 2024, the lineage distribution of SARS-CoV-2 in England exhibited significant genetic shifts, as illustrated by the prevalence trajectories of dominant lineages during this period (Figure 7C) sampled from GISAID [83]. The JN.1 lineage, including sub-lineages JN.1.1 and JN.1.4, experienced rapid expansion, attributed to distinct molecular adaptations [84]. Using e3SIM, we simulated these evolutionary dynamics by integrating genetic variation into a realistic contact network model. We constructed a contact network with 12,907 nodes based on the Haslemere dataset [85, 86], a publicly available social interactions dataset used to study COVID-19 control strategies [21], assuming a power-law degree distribution. We defined a genetic architecture where JN.1 has higher transmissibility than BA.2.86.1 and EG.5.1 [84] and randomly seeded the host population with sequences from GISAID (Figure 7A). Samples were collected sequentially during the simulation. The lineage composition trajectories in e3SIM’s simulation (Figure 7B) shows the increasing prevalence of the JN.1 lineage over time, with declines in the BA.2.86.1 and EG.5.1 lineages. This pattern closely mirrors the observed lineage frequencies in real data (Figure 7C). Quantitative discrepancies between the simulated and empirical data may be attributed to the real contact network not strictly following a power-law degree distribution, genetic traits other than transmissibility, and environmental factors the impact pathogen dynamics.

**Figure 7.**
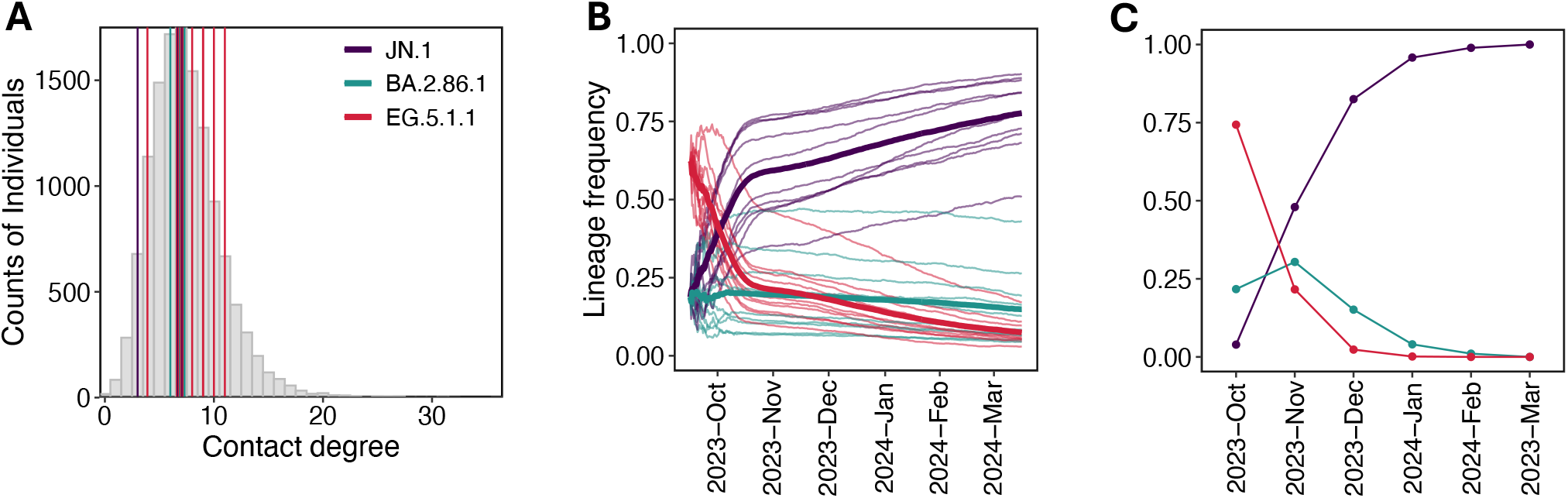
Comparison of simulated SARS-CoV-2 lineage distribution with observed data. (A) Degree distribution of the contact network used for simulating the Haslemere SARS-CoV-2 outbreak. Vertical lines indicate the contact degrees of hosts seeded by the initial SARS-CoV-2 sequences. (B) Simulated lineage frequency trajectories for sampled JN.1, BA.2.86.1, and EG.5.1.1 over time across ten simulation replicates. Bold lines represent the average trajectory across all replicates. (C) Observed lineage frequency trajectories for JN.1, BA.2.86.1, and EG.5.1.1 over time in the United Kingdom, aggregated monthly.

#### 3.1.4 Mtb transmission dynamics in Karonga, Malawi: impact of contact networks

The Karonga Prevention Study collected over 2,000 *Mtb* whole-genome sequences from the Karonga district in northern Malawi, a region with over 300,000 inhabitants and a high tuberculosis incidence (*∼* 100 cases per 100,000 population), between 1995 and 2014 [87–89]. To evaluate the impact of host population structure on the Karonga epidemic, we compared two network models: the Erdős–Rényi network, a null model without population structure, and the Barabási–Albert networks, which incorporates population structure with hubs. Each network comprised 300,000 nodes, corresponding to the population size of Karonga [89]. We simulated the Karonga epidemic over a 20-year period by seeding the host population with all the sampled sequences from the Karonga dataset between 1995 and 1998 (*n* = 375), reflecting the incidence rate of approximately 100 cases per 100,000 population [89]. Genetic and timed trees were inferred from sequential sampling conducted during the final three years. The internal branch length distributions derived from the genetic trees were stable across replicates (Figure S8), with the distribution from the Erdős–Rényi network closely matching those inferred from real data (Figure 8D). The past effective population size trajectories inferred from the timed trees of the Erdős–Rényi model (Figure 8B) using a Bayesian nonparametric phylodynamic method [90] aligned well with the trajectory inferred from real data (Figure 8A). In contrast, trajectories from the Barabási–Albert model (Figure 8C) deviated more significantly from the past effective population size trajectory inferred from the real data. These results suggest that an Erdős–Rényi network may adequately represent the actual host contact structure during the Karonga *Mtb* epidemic [91]. Our simulation framework shows close alignment with observed epidemiological and genetic patterns, indicating its effectiveness in modeling real-world epidemics. While a fully rigorous model selection process would require more comprehensive matching of simulation outputs to empirical data using advanced statistical techniques, our current analyses provide robust qualitative validation. These results highlight that our simulation framework is a valuable tool for studying epidemiological dynamics and assisting in model selection.

**Figure 8.**
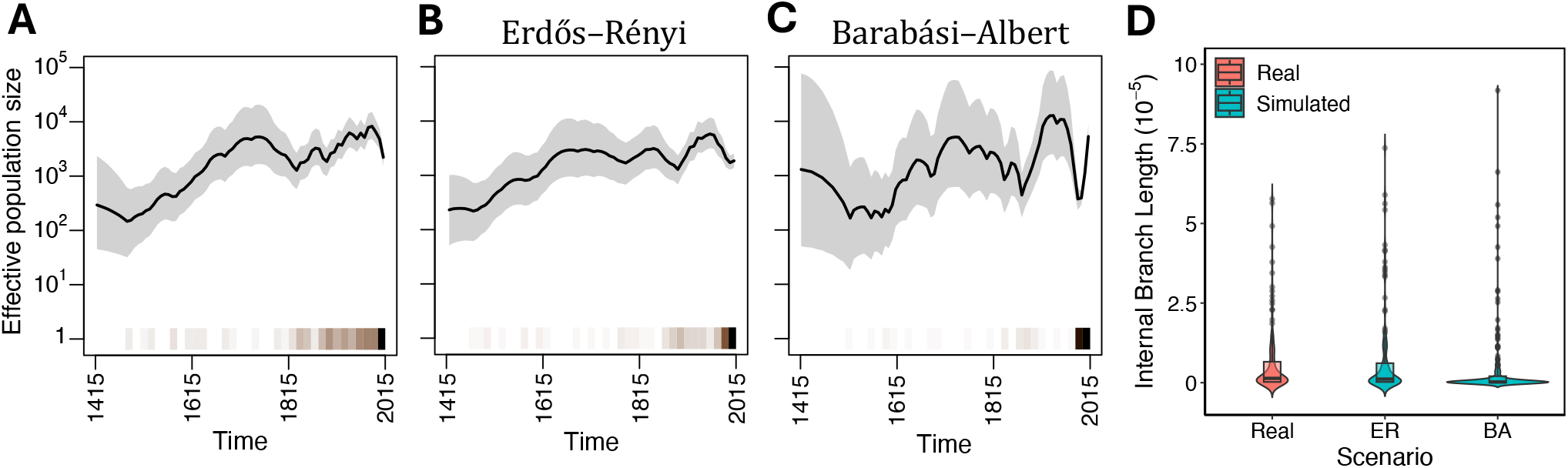
Comparison of inferred effective population size histories and internal branch length distributions between the simulated and real Karonga datasets. (A) Inferred effective population size trajectory for the real dataset. (B) Inferred effective population size trajectory for the simulated dataset using Erdős–Rényi network. (C) Inferred effective population size trajectory for the simulated dataset using Barabási–Albert network. The heatmap at the bottom indicates the density of coalescent events across all plots; the darker the color, the greater the number of events occurring within a given time interval. (D) Violin plot showing the internal branch length distribution of the genetic tree inferred from samples in one simulation replicate, compared to the distribution inferred from real data. The results corresponding to each scenario across all six replicates can be found in Figures S6–S8.

### 3.2 Runtime profiling

The runtime of `OutbreakSimulator` module is influenced by various configuration settings, such as the number of seeds, genome length, number of traits, mutation rate, and the expected outbreak size determined by all the epidemiological parameters. We conducted runtime profiling, focusing on key parameters, including host population size, sample size and base transmission probability *β*. Simulations were conducted on a MacBook Pro equipped with an Apple M2 Pro CPU and 32GB of RAM, running macOS Sonoma version 14.5. Runtime vary significantly with host population sizes (Figure 9). For a host population of 100,000, `OutbreakSimulator` completed the simulation in 16 minutes, with *∼* 60% of the host population being in infected state once the equilibrium was reached after *∼* 100 ticks. For smaller host populations, an increase in sample size per tick markedly increased runtime. For example, with a host population of 10,000, increasing the expected maximum sample size per tick from 1 to 10 resulted in a 3.88-fold increase in average runtime. Conversely, for larger host populations, the runtime remained relatively stable, increasing by only about 4% when the expected maximum sample size per tick was adjusted from 1 to 10 for a host population of 100,000. The base transmission probability (*β*) significantly influences the outbreak sizes, subsequently impacting runtime. We performed simulations on a Linux system using the simulation parameter set detailed in Figure 9B, varying only the base transmission probability to generate different outbreak sizes. Figure S9 illustrates the relationship between transmission probability and runtime.

**Figure 9.**
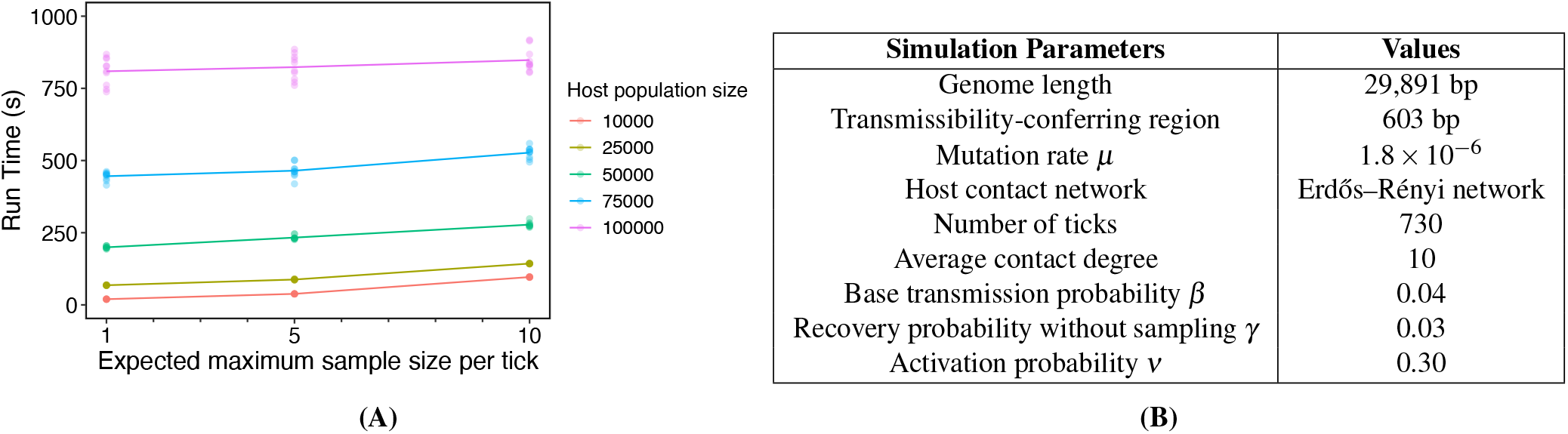
Runtime profiling on a personal computer using a single core. (A) Dots represent individual run times, while lines indicate the average run time across ten replicates for each parameter set. The *x*-axis represents the expected maximum sample size per tick, calculated as the product of the host population size and the sampling probability per tick. Sequential sampling probability is adjusted to keep the same maximum expected sample size (1, 5, 10) for each host population size. This ensures a consistent maximum expected sample size per tick across different host population sizes by adjusting the sequential sampling probability *ε*_*s*_ (Table S4). (B) Simulation parameters used for the runtime profiling.

## 4 Discussion

We introduced e3SIM, an epi-eco-evo simulation framework specifically designed for studying how the eco-evolutionary dynamics of pathogens impact disease transmission. With its modular and scalable architecture, e3SIM supports a broad range of epidemiological and population-genetic complexities, realistically integrating epidemiological compartmental models with host-population contact networks and quantitative-trait models for pathogens. Additionally, e3SIM features a user-friendly graphical interface, enhancing accessibility for users from diverse backgrounds. Our simulation examples, including scenarios with SARS-CoV-2 and *Mtb*, demonstrated e3SIM’s advanced capabilities in concurrently modeling disease dynamics and molecular evolution while incorporating environmental factors.

Several methods (Table S5) have attempted to model aspects of epi-eco-evo processes. However, these approaches often fail to fully integrate with epidemiological compartmental models, neglect host contact heterogeneity and its impact on disease dynamics, or overlook time-varying environmental effects. While Cárdenas et al. [35] introduced several advanced features, their method was limited to the SIRD compartmental framework, excluding important aspects such as latency for pathogens like *Mtb*, and did not incorporate molecular evolution models. Moreover, their approach assumed uniform contact rates within structured populations, restricting its flexibility to model individual-level interactions effectively. Another significant limitation of Cárdenas et al. [35] is its inability to generate a true genealogy of pathogens, an essential component for phylodynamics studies of genomic epidemiology [92–95]. Additionally, the complexity of the simulator and the absence of a user-friendly interface further restrict its practical application. In contrast, e3SIM addresses these limitations by offering comprehensive functionalities for modeling epi-eco-evo processes while maintaining ease of use, establishing it as a powerful and versatile tool in genomic epidemiology.

While e3SIM is a powerful tool, it has several limitations. First, it does not include recombination, a crucial evolutionary mechanism in many pathogens [96], such as HIV [97] and hepatitis C [98]. This limitation could restrict its ability to fully capture the genetic diversity and evolutionary dynamics of non-clonally reproducing pathogens. Additionally, e3SIM does not model host-pathogen co-evolution, an important factor driving significant evolutionary changes in both host and pathogen populations [99, 100]. However, both limitations can be readily addressed by incorporating recombination and multispecies modeling features of SLiM 4 [42]. Further, the current implementation employs a static contact network, which does not reflect the dynamic nature of human interactions. In real-world scenarios, contact patterns can change over time due to factors such as social behaviors [101, 102], public health interventions [103, 104], and seasonal variations [105, 106]. While dynamic contact networks have been used in epidemiology studies, no current genomic epidemiology simulators explicitly model these networks to our knowledge. Future enhancements of e3SIM will incorporate dynamic contact networks to model changes in host contact patterns more accurately. This will involve integrating time-varying network models and stochastic processes to simulate the formation and dissolution of contacts [107, 108].

Despite these limitations, e3SIM represents a significant advancement in the field of genomic epidemiology, providing a robust and flexible platform for simulating the complex interplay of epidemiological, ecological, and evolutionary processes. Its modular design ensures that it can be easily extended to address current limitations and incorporate future advances, making it a valuable tool for researchers and public health professionals to understand and mitigate the spread of infectious diseases.

## 5 Methods

### 5.1 Normalization in GeneticEffectGenerator

Determining an appropriate range for effect sizes can be challenging when using the random generation mode. e3SIM includes an optional feature that scales effect sizes based on the total number of simulation ticks (*g*) and the mutation rate per site per tick (*μ*). This normalization heuristically adjusts the expected trait value at the simulation’s conclusion (Section 2.1.3) according to the user-specified trait value *T*. Specifically, the normalized effect sizes 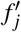 for mutations in each causal genomic element *j* and each trait are calculated as:

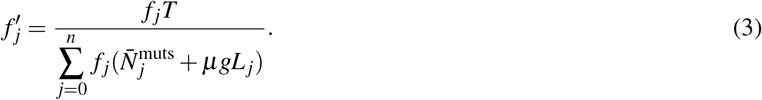

Here, *f* _*j*_ represents the unnormalized effect sizes for each mutation in genomic element *j*. The term 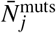 denotes the average number of mutations in genomic element *j* across all seeds, as defined by `GeneticEffectGenerator`. *L*_*j*_ denotes the length of genomic element *j*, and *g* is the total number of ticks in the simulation. The denominator represents the expected trait value at the final tick of `OutbreakSimulator` before normalization. This normalization provides a baseline that users can manually adjust according to their specific needs.

### 5.2 Epi-eco-evo dynamics in fast-evolving pathogens (Section 3.1.1)

We simulated a SARS-CoV-2 outbreak in a population of 10,000 individuals over 720 ticks, divided into three epochs, with each tick representing one day. The contact network was generated using the Barabási–Albert model in `NetworkGenerator` (`-model BA -m 2`). We defined a genetic architecture comprising one transmissibility trait and two drug resistance traits. The drug resistance traits were introduced in the second and third epochs, respectively, to simulate resistance to two different drug treatments. The transmissibility trait was governed by a 351 bp locus (positions 650–1000), with mutations within this locus affecting transmissibility. The transmissibility trait locus had an effect size of 0.2. The first drug resistance trait was governed by a 101 bp locus (positions 450–550), while the second was governed by a 301 bp locus (positions 4300-4600). Each drug resistance trait locus had an effect size of 0.3. The host population was seeded with a single SARS-CoV-2 reference genome. The first and second treatment stages began at 250 and 500 ticks, respectively. Samples were sequentially collected with a probability of 0.0001 per tick across all ten replicates. The complete configuration for `OutbreakSimulator` is detailed in Configuration File S1.

### 5.3 Epidemic dynamics with superspreaders (Section 3.1.2)

We simulated a *Mtb* outbreak in a host population of 10,000 individuals over a 10-year period using e3SIM. The population structure was modeled with two network types: 1) an Erdős–R’enyi network (`-p_ER 0.001`) to represent a homogeneous network, and 2) a Barabási–Albert network (`-m 2`) to account for superspreaders, defined as hosts with a higher number of contacts compared to the majority of others. Seed sequences for the simulation were generated with the `SeedGenerator` module using the Wright-Fisher model and the *Mtb* H37Rv reference genome (GCF_000195955.2) [51]. The genetic architecture included five loci for transmissibility, randomly generated from the reference genome’s `GFF` file from NCBI using `GeneticEffectGenerator` with the following parameters:

~~~
-es_low 1 -es_high 5 -causal_size_each 5 -normalize T -sim_generation 500 -mut_rate 4.4e-9.
~~~

Each trait locus had an effect size between 0.10 and 0.37 (see Code Availability). Using the two networks generated by `NetworkGenerator`, we simulated three different scenarios to demonstrate the effect of superspreaders and contact networks on epidemic dynamics. For each scenario, samples were collected sequentially with a probability of 0.00002 per tick across all six replicates.

In the first scenario, we randomly assigned the five seeds sequences to hosts within the Erdős–Rényi networks (Figure 6A) using the completely random mode of `SeedHostMatcher`. In the second scenario, high-transmissibility seeds were matched to socially active hosts within the Barabási–Albert network, while low-transmissibility seeds were matched to less socially active hosts (Figure 6B). The `HostSeedMatcher` parameters were:

~~~
-match_scheme ‘{“0”: “Ranking”, “1”: “Ranking”, “2”: “Ranking”, “3”:”Ranking”}’
-match_scheme_param ‘{“0”: 600, “1”: 700, “2”: 4000, “3”: 5000}’.
~~~

In the third scenario, low-transmissibility seeds were matched to socially active hosts within the Barabási–Albert network, while high-transmissibility seeds were matched to less socially active hosts. The `HostSeedMatcher` parameters were set to:

~~~
-match_scheme ‘{“0”: “Ranking”, “1”: “Ranking”, “2”: “Ranking”, “3”: “Ranking”}’
-match_scheme_param ‘{“0”: 7000, “1”: 8000, “2”: 300, “3”: 500}’.
~~~

The full configuration for `OutbreakSimulator` can be found in Configuration File S2.

### 5.4 SARS-CoV-2 epidemic dynamics in Haslemere (Section 3.1.3)

#### 5.4.1 Contact network generation

We utilized the Haslemere dataset, a publicly available resource for modeling infectious disease dynamics at the population level [85, 86]. This dataset, collected in Haslemere, England—a town with a population of 12,907 according to the 2021 national census [109]—was previously used to study COVID-19 control strategies [21]. It includes pairwise distances between users of the BBC Pandemic Haslemere app over a three-day period.

From the original contact tracing data provided by Kissler et al. [85] in `CSV` format, we filtered the data to define a valid contact as any instance where two participants were within 10 meters of each other, adopting the approach of Firth et al. [21]. Based on this filtered contact information, we constructed a contact matrix. COVID-19 transmission has often been shown to exhibit power-law characteristics due to the presence of superspreaders and heterogeneous contact patterns [110–112]. We therefore modeled the Haslemere contact network degree distribution as a power law. We calculated the degree density of this contact network (*n*_node_ = 469, *n*_edge_ = 1753) and fitted it to a power-law distribution using the `displ$new` function in the `poweRlaw` package [113, 114] in R. The estimated exponent was *γ* = 6.21 for *x*_*min*_ ≥ 19 according to the best goodness-of-fit.

To scale the network to the size of the Haslemere population, we defined the number of nodes as *n*_node_ = 12, 907 and scaled the total number of contacts while maintaining the same average contact degree as the original network 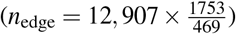. We generated the larger network using the `sample_fitness_pl` function in `igraph` [115], specifying the number of nodes, edges, and the exponent of the power-law degree distribution:

~~~
exponent.in = -1, loops = FALSE, multiple = FALSE, finite.size.correction = TRUE.
~~~

The network was stored in adjacency list format and processed by `NetworkGenerator` to produce a contact network file for simulation.

#### 5.4.2 Processing of SARS-CoV-2 genetic data

We identified the top five SARS-CoV-2 lineages in England from October 2023 to March 2024 according to GISAID [116]: JN.1, JN.1.4, BA.2.86.1, JN.1.1, and EG.5.1.1. Complete high-quality sequences of these lineages, as defined by GISAID, were downloaded in `FASTA` format. These sequences were aligned to the reference genome hCoV-19/Wuhan/WIV04/2019 (WIV04) using MAFFT v7.522 [117] with the parameters ’`-6merpair -keeplength -addfragments`’. The aligned sequences were converted to `VCF` format using custom R scripts (see Code Availability). Ambiguous alleles (e.g., Y, K) were manually resolved to either the reference allele or the major alternative alleles of other samples, retaining only A, C, G, and T alleles. Based on the initial lineage composition in October 2023 recorded in GISAID [83], we selected the earliest 3 samples from JN.1, 1 sample from JN.1.4, 3 samples from BA.2.86.1, 1 sample from JN.1.1, and 10 samples from EG.5.1.1 as seeding sequences for our simulation.

Using the whole-genome sequences of all samples from the top five lineages collected between October 2023 and March 2024 in England (1,143 JN.1 samples; 228 JN.1.4 samples; 181 BA.2.86.1 samples; 156 JN.1.4 samples; 157 EG.5.1.1 samples), we inferred the substitution rate matrix **Q** based on genetic distance (substitutions/site) using IQ-TREE 2 v2.2.2.7 [68] with the GTR model (`-m GTR`). Since the estimated **Q** is in the time unit of the expected time for a single mutation, we rescaled **Q** in the unit of days using the molecular clock rate of *μ* = 9 × 10^−7^ substitutions/site/day for SARS-CoV-2 [118]. We then used the `scipy.linalg.expm` function from SciPy package [119] in Python to compute the transition probability matrix **P** = *e*^**Q***t*^ for one tick, assuming one tick equals one day, which was subsequently used in the e3SIM simulation.

#### 5.4.3 e3SIM simulation

The BA.2.86 and JN.1 lineages possess mutations that confer higher growth rates and potential for increased transmissibility [84, 120]. In contrast, the success of EG.5.1 is attributed to factors other than increased transmissibility, making it a less critical focus for transmission dynamics [121]. Building on these observations, we used the Pangolin tool [122] to extract lineage-defining mutations for JN.1 and BA.2.86, assigning transmissibility genetic architecture to a subset of them [122]. The K1973R mutation in orf1a, present in both JN.1 and BA.2.86, was assigned an effect size of 0.4. Based on the rapid growth of the JN.1 lineage during the study period (Figure 7C), we assigned an effect size of 0.35 to the R3821K mutation, which is exclusive to JN.1. These sequences were randomly seeded into the generated contact network (Figure 7A) and sampled sequentially with a probability of 0.0007 per tick throughout the simulation. The full configuration for `OutbreakSimulator` is detailed in Configuration File S3. For each of the ten simulation replicates, the lineage composition of samples was computed, with JN.1, JN.1.4, and JN.1.1 collapsed into JN.1 (Figure 7B). These resulting lineage distributions were then compared to GISAID data collected from October 2023 to March 2024, aggregated by month (Figure 7C).

### 5.5 Mtb transmission dynamics in Karonga (Section 3.1.4)

#### 5.5.1 Contact network generation

We employed two types of contact networks for simulations in Karonga: Erdős–Rényi and Barabási–Albert networks. The Erdős–Rényi network was generated using the `NetworkGenerator` in e3SIM with the parameter `-p_ER 0.00001`. The Barabási–Albert network was also generated using the `NetworkGenerator` in e3SIM with the parameter `-m 2`.

#### 5.5.2 Processing of Mtb genetic data

We processed the concatenated SNP files from *Mtb* 1,857 samples as described in Sobkowiak et al. [89], using custom Python scripts (see Code Availability) to map them to the *Mtb* H37Rv reference genome (GCF_000195955.2) [51], generating whole-genome sequences. IQ-TREE 2 v2.2.2.7 [68] with the GTR model (`-m GTR`) was then used to infer the substitution rate matrix **Q** based on genetic distance (substitutions/site). Using the pipeline in Section 5.4.2, we computed the transition probability matrix **P** for one tick from **Q**, with the molecular clock rate of *μ* = 1.133393 × 10^−7^ substitutions/site/year for *Mtb* (0.5 substitutions per genome per year) [123], and one tick representing three days.

#### 5.5.3 e3SIM simulation

We initiated the simulation with all sampled sequences from 1995–1998 (*n* = 375) and simulated a 20-year period across six replicates per each contact network. Sampling occurred sequentially only in the last three years. In cases where multiple samples were taken from the same host, only the first sampled genome per host was retained. We performed a uniform sub-sampling to match the final sample size of the Karonga dataset from 2012-2014 (*n* = 168) [88, 89]. No genetic architecture was invoked for any trait during the simulation. The full configuration for `OutbreakSimulator` is detailed in Configuration File S4.

#### 5.5.4 Internal branch length distribution and effective population size inference

We converted logged sampling times in ticks to real-time scale and transformed the `VCF` file into `FASTA` format by mapping SNPs to the H37Rv genome [124] using a custom R script (see Code Availability). We used IQ-TREE 2 v2.2.2.7 [68] with the GTR model (`-m GTR`) on sequences sampled from both the simulation and the 2012–2014 Karonga dataset [89]. From these genetic trees, we calculated branch length distributions. To infer the timed tree, we rescaled each branch length of the genetic tree to represent the number of substitutions across the entire genome. We ran the `bactdate` function from the BactDating package [125] in R on the rescaled tree with the following parameters:

~~~
initMu = 0.5, nbIts = 1e6, thin = 1e3, updateMu = F,
updateRoot = T, model = ‘strictgammaR’, useCoalPrior = T.
~~~

From the resulting timed tree, we inferred the past effective population size trajectory via Bayesian nonparametric phylodynamic reconstruction using the `BNPR` method in the phylodyn package [90] with default settings.

### 5.6 Runtime profiling (Section 3.2)

To benchmark the performance of the `OutbreakSimulator` module in e3SIM, we conducted tests on a MacBook Pro equipped with an Apple M2 Pro CPU and 32GB of RAM, running macOS Sonoma version 14.5. The `NetworkGenerator` was configured to produce Erdős–Rényi networks with an average degree of 10, varying the host population size from 10,000 to 100,000 (`-p_ER` from 0.001 to 0.0001). The population sizes tested were 10,000, 25,000, 50,000, 75,000, and 100,000. In each network, the host with the median contact degree was seeded with the SARS-CoV-2 reference genome, and samples were collected throughout the simulation. Sampling probabilities were adjusted to achieve maximum sample sizes of 1, 5, or 10 per tick for each population size, and runtimes were recorded (Figure 9). We conducted additional performance benchmarks on a Linux system to evaluate runtime across various outbreak sizes, using a host population of 100,000 and an Erdős–Rényi contact network. We kept the same parameters as those listed for the 100,000 host population in Figure 9B, except for varying the base transmission probability (*β*) from 0.01 to 0.06. Sequential sampling was performed with a probability of 0.0001 per tick. The Linux benchmark experiments utilized a single CPU and 5 GB of memory.

## 6 Data Availability

The reference genome for all SARS-CoV-2 simulations in this study is the official reference genome employed by GISAID: hCoV-19/Wuhan/WIV04/2019 (WIV04). The SARS-CoV-2 sequencing data used in the Haslemere simulation are based on metadata associated with 1,865 sequences collected in England between October 1, 2023, and March 24, 2024. These sequences are available on GISAID as of April 2, 2024, via doi:10.55876/gis8.240614pc.

The H37Rv reference genome and its `GFF` annotation file, used for all *Mtb* simulations in this study, are accessible via NCBI GCF_000195955.2. The *Mtb* sequencing data for the Karonga simulations is from Sobkowiak et al. [89].

## 7 Code Availability

Our open-source software, e3SIM, is available on GitHub at EpiEvoSoftware/original_pipeline. The repository also includes a comprehensive manual.

The code used for analyzing both real and simulated data, as well as generating the plots presented in this manuscript, has been uploaded to Zenodo: doi:10.5281zenodo.11715270.

## Acknowledgements

PX, SL, AH, VZ, WTL, BS, MHC, CC, TC, KYR, MTW, AGC, and JK were supported by the National Institutes of Health Grant P01AI159402. BCH and PWM were supported by the National Institutes of Health under awards R01HG012473 and R35GM152242.

