## Supplementary Information for "e3SIM: epidemiological-ecological-evolutionary simulation framework for genomic epidemiology"

**Table S1.** Modules of e3SIM.

| | Module | Input in random generation mode | Output | Function |
|---|---|---|---|---|
| Pre-simulation modules | **`NetworkGenerator`** **(Section 2.1.1)** | • Specification of a random network model and parameters. | • Adjacency list specifying contact structure. | Generate a host contact network. |
| Pre-simulation modules | **`SeedGenerator`** **(Section 2.1.2)** | • Pathogen reference genome.<br>• Models and parameters.<br>• Optional: Contact network from `NetworkGenerator`. | • `VCF` files for each seed sequence.<br>• `NWK` file of seed phylogeny. | Generate sequences for pathogen seeds. |
| Pre-simulation modules | **`GeneticEffectGenerator`** **(Section 2.1.3)** | • `GFF` file for the reference genome.<br>• Parameters for genetic architecture.<br>• Optional: Seed sequences from `SeedGenerator` | • `CSV` file detailing genetic architecture.<br>• `CSV` file of trait values for all seeds based on genetic architecture generated by `SeedGenerator`. | Create a genetic architecture for the pathogen genome to be used by `OutbreakSimulator` for trait value calculation. |
| Pre-simulation modules | **`HostSeedMatcher`** **(Section 2.1.4)** | • Host contact network from `NetworkGenerator`.<br>• Matching scheme and parameters. | • `CSV` file specifying host-seed assignments. | Match each seed to a host based on user-defined criteria. |
| Main & Post-simulation modules | **`OutbreakSimulator`** **(Section 2.2)** | • Host contact network produced by `NetworkGenerator`.<br>• Seeds' sequences produced by `SeedGenerator`.<br>• Genetic architecture produced by `GeneticEffectGenerator`.<br>• Seed-host matching file produced by `HostSeedMatcher`.<br>• Pathogen reference genome.<br>• Simulation configuration file in JSON format. | • `VCF` files of sampled pathogen genomes.<br>• Log file documenting all the epidemiological events.<br>• `NWK` file of the genealogy of the sampled genomes.<br>• SEIR trajectory plot.<br>• Lineage trajectory plot.<br>• Plotted genealogy of the sampled pathogens.<br>• Metadata file of the genealogy of sampled pathogens. | Execute the whole simulation based on the provided configuration and input files, and process the resulting output data. |

**Table S2.** Modules of `OutbreakSimulator`.

| Module | Relevant configurations (in the configuration file) | Function |
|---|---|---|
| **`Reproduction`** **(Section 2.2.3)** | `genetic_architecture`, `EvolutionModel` | Defines between-host transmission and within-host reproduction events. Each entry in `genetic_architecture` is a list of integers, with each integer corresponding to an epoch as specified in `epoch_changing`, indicating the genetic architecture used during each epoch. |
| **`Compartmental model`** **(Section 2.2.4)** | `transiton_prob`, `model`, `genetic_architecture` | Defines the compartmental model for the simulation, specifying the transition probabilities between compartments S (Susceptible), E (Exposed), I (Infected), and R (Recovered). These probabilities are applied to each pathogen or host per time tick. Each probability entry must be a list corresponding to the number of epochs specified in `epoch_changing`. For transmission and recovery events, the actual probability is also influenced by the genetic architecture for pathogen transmissibility and drug resistance. |
| **`Sampling`** **(Section 2.2.5)** | `transiton_prob`, `massive_sampling` | Specifies the sampling method during the simulation. Sequential sampling events are included in `transiton_rate`, while concerted sampling events are defined separately for each tick when they occur. |

**Table S3.** List of symbols for epidemiological parameters, visualized in Figure 3C. All the probabilities on a per-tick basis.

| Symbol | Description | Target level |
|---|---|---|
| $\omega$ | Probability of immunity loss for a recovered host | Host |
| $\beta$ | Base probability of successful infection event for one effective contact | Host |
| $\zeta$ | Probability of latent infection for each successful infection event | Host |
| $\phi$ | Probability of deactivation for an infected host | Host |
| $\nu$ | Probability of activation for an exposed host | Host |
| $\tau$ | Probability of transitioning to recovered state for an exposed host | Host |
| $\gamma$ | Base probability of clearance for a pathogen residing in an infected host | Pathogen |
| $\varepsilon_s$ | Probability of being sampled in a sequential sampling event for an infected host | Host |
| $\delta_s$ | Probability of recovery for an infected host following a sequential sampling event | Host |
| $\varepsilon_c$ | Probability of being sampled in a concerted sampling event for an infected host | Host |
| $\delta_c$ | Probability of recovery for an infected host following a concerted sampling event. | Host |

**Table S4. Sampling probability per tick for each active pathogen used in runtime profiling.** The host size refers to the number of hosts in the population. The expected maximum sample size per tick, calculated as Host size × Sampling probability, represents the expected number of pathogens sampled per tick if all hosts are infected by a single pathogen. The actual sample size per tick during the simulation is generally smaller than this expected maximum, depending on the outbreak size. The runtime profiling results and other parameters used are described in Section 3.2

| | | **Expected (maximum) sample size each tick** | | |
|---|---|---|---|---|
| | | **1** | **5** | **10** |
| **Host size** | **10000** | 0.0001 | 0.0005 | 0.001 |
| | **25000** | 0.00004 | 0.0002 | 0.0004 |
| | **50000** | 0.00002 | 0.0001 | 0.0002 |
| | **75000** | 0.000013 | 0.000067 | 0.00013 |
| | **100000** | 0.00001 | 0.00005 | 0.0001 |

**Table S5. Comparison of e3SIM with other simulators for genomic epidemiology.**

| Simulator Name | Compartmental Model | Epi-Eco-Evo Coupling | Host Population | Within-Host Reproduction | Time-Varying Parameters | Recombination | Host Genomic Effects |
|---|---|---|---|---|---|---|---|
| **e3SIM [this paper]** | **SEIRS User-defined** | **True** | **Contact network** | **True** | **True** | **False** | **False** |
| FAVITES [1] | GEMF model [2] | False | Contact network | False | False | False | False |
| nosoi [3] | SIRD | False | Structured | False | False | False | False |
| opqua [4] | SIRD | True | Structured | True | True | True | False |
| outbreaker2 [5] | SI | False | Contact network | True | True | False | False |
| SANTA-SIM [6] | – | True | – | False | True | True | False |
| SEEDY [7] | SIR | False | Contact network | True | False | False | False |
| TiPS [8] | User-defined | False | Structured | False | False | False | False |
| Vgsim [9] | SI | True | Structured | False | True | False | False |
| Rasmussen et al. [10] | – | True | – | False | False | False | False |

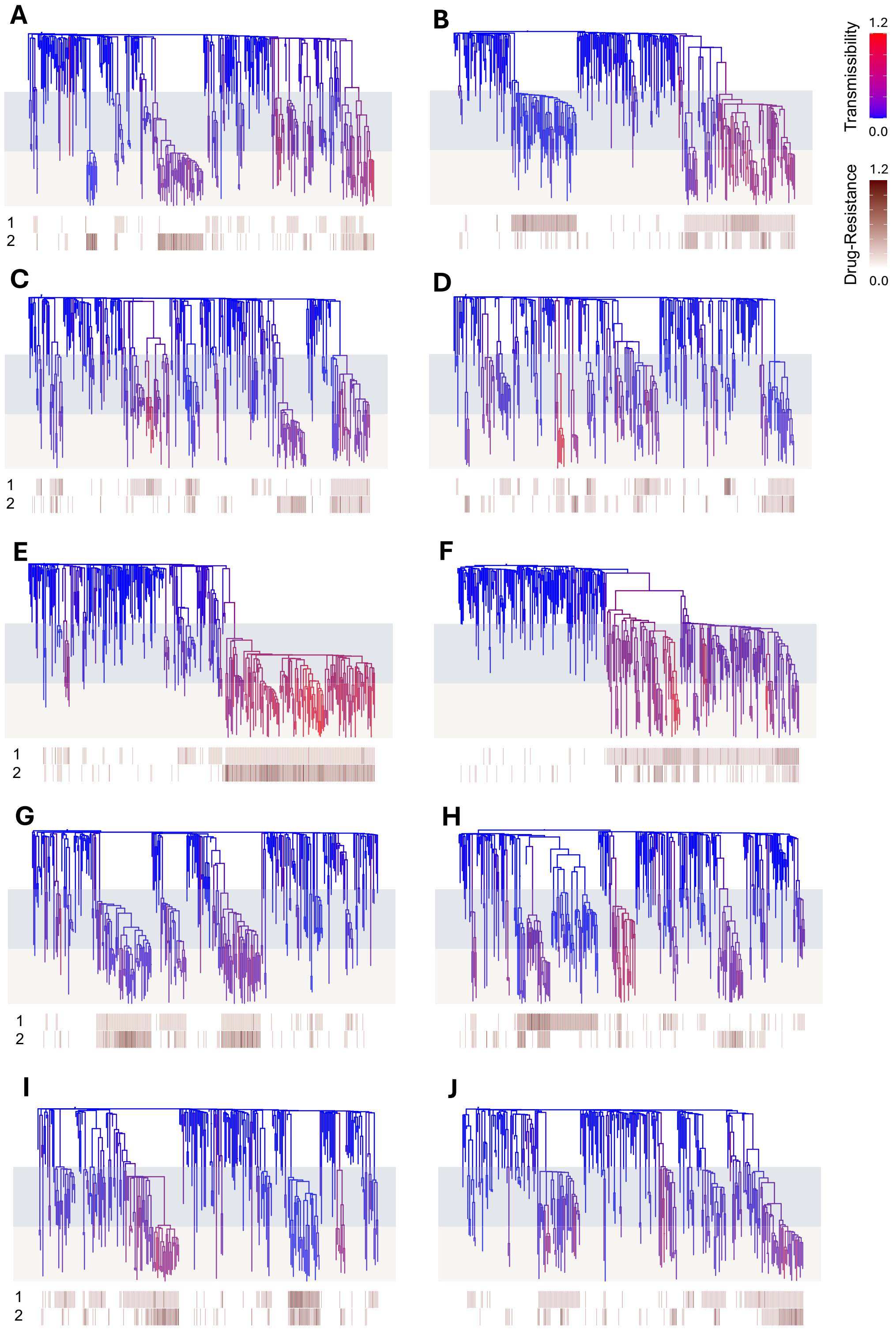


**Figure S1. Genealogies of sampled pathogens illustrating the emergence of new variants and epi-eco-evo dynamics in SARS-CoV-2 across ten simulation replicates.** Tree branches are colored by relative transmissibility (blue: lowest, red: highest), with a heatmap indicating drug-resistance trait values for each pathogen genome (tips). Heatmap rows represent the first and second drug-resistance traits. The shaded backgrounds denote treatment stages 1 and 2. Figure A (the first replicate) is presented in Figure 5A. Across all replicates, lineages with higher transmissibility and drug resistance gained higher prevalence in a drug treatment environment. Simulation details are provided in Section 5.3, supplementing Section 3.1.1.

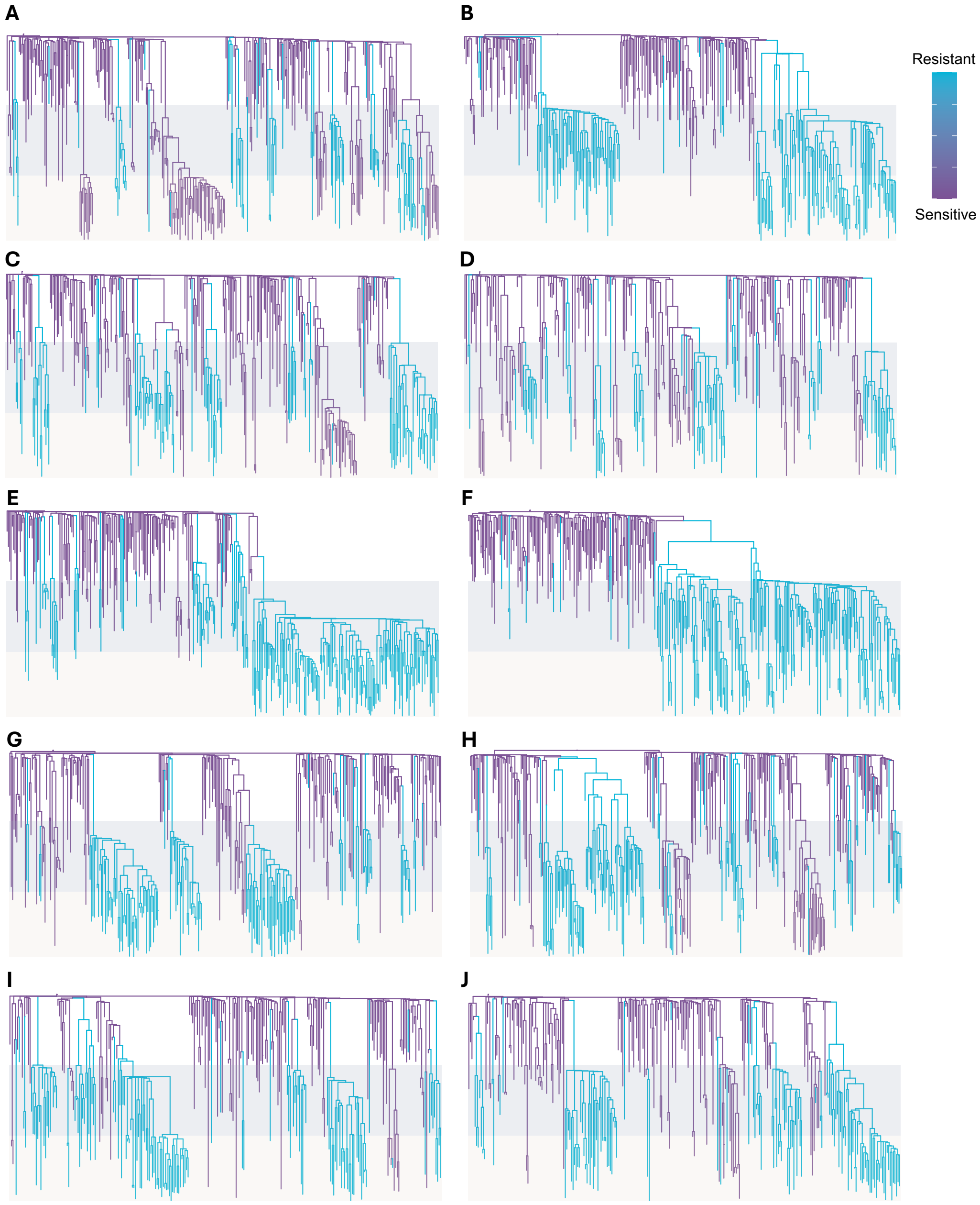


**Figure S2. Genealogies of sampled pathogens illustrating the emergence of drug resistance mutations and epi-eco-evo dynamics in SARS-CoV-2 across ten simulation replicates.** These trees are identical to those in Figure S1, but branches are colored by resistance to the first treatment (purple: sensitive, light blue: resistance). The figure design follows Figure S1. Simulation details are provided in Section 5.3, supplementing Section 3.1.1.

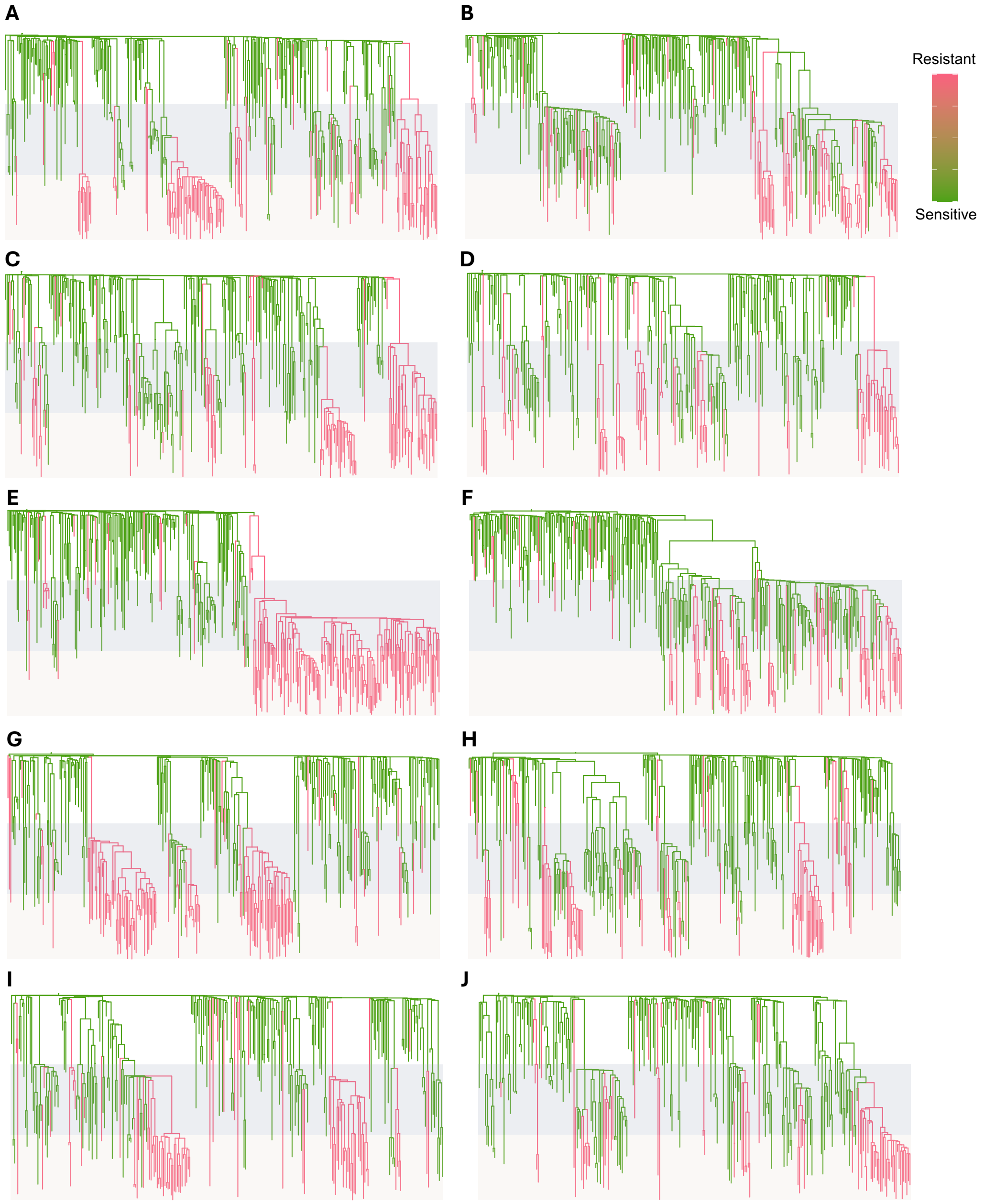


**Figure S3. Genealogies of sampled pathogens illustrating the emergence of drug resistance mutations and epi-eco-evo dynamics in SARS-CoV-2 across ten simulation replicates.** These trees are identical to those in Figure S1, but branches are colored by resistance to the second treatment (green: sensitive, pink: resistance). The figure design follows Figure S1. Simulation details are provided in Section 5.3, supplementing Section 3.1.1.

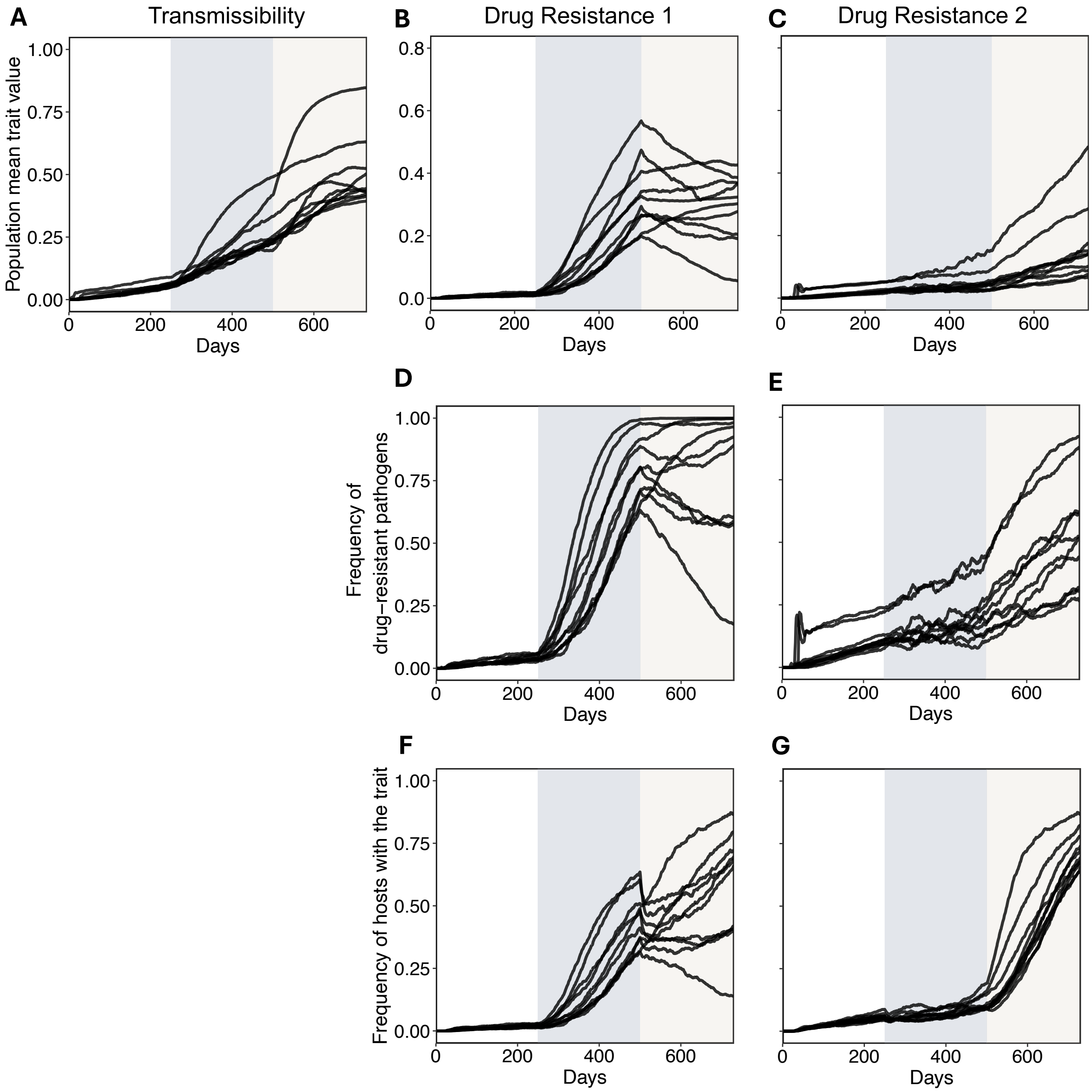


**Figure S4. Population mean trait values and the frequency of hosts with the trait over time across ten simulation replicates.** (A) Mean transmissibility. (B) Mean resistance to the first drug. (C) Mean resistance to the second drug. (D) Frequencies of pathogens with resistance to the first drug. (E) Frequencies of pathogens with resistance to the second drug. (F) Frequencies of hosts carrying a pathogen resistant to the first drug. (G) Frequencies of hosts carrying a pathogen resistant to the second drug. Mean values and frequencies were calculated for all pathogens and hosts in the population at each time point. Shaded regions indicate treatment stages 1 and 2. Detailed simulation parameters are in Section 5.3, supplementing Section 3.1.1.

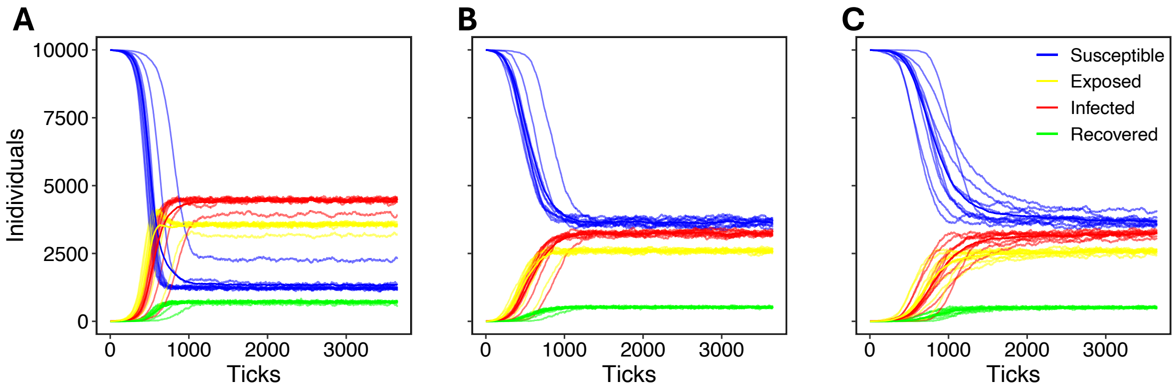


**Figure S5. Effect of superspreaders and contact network on *Mtb* epidemic dynamics.** Each curve represents one simulation run, with ten replicates combined in a single plot for each scenario. Simulation details are in Section 5.4, supplementing Section 3.1.2. (A) SEIR trajectory for the Erdős–Rényi network using random seed matching (Figure 6A). (B) SEIR trajectory for the Barabási–Albert network, with highly transmissible seeds assigned to highly connected hosts, and less transmissible seeds to less connected hosts (Figure 6B). Matching scheme detailed in Section 5.4. (C) SEIR trajectory for the Barabási–Albert network, with highly transmissible seeds assigned to less connected hosts, and less transmissible seeds to highly connected hosts (Figure 6C). Matching scheme detailed in Section 5.4.

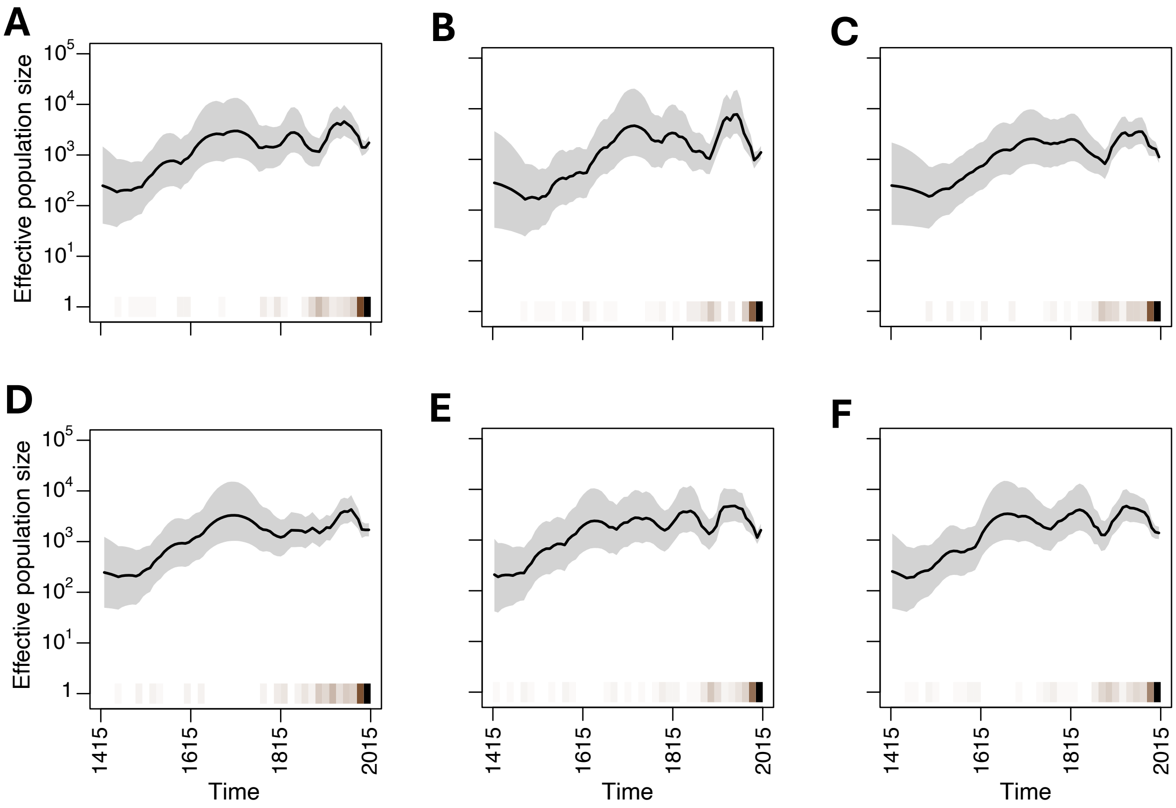


**Figure S6. Effective population size trajectory inferred by the BNPR method for the Karonga simulation using the Erdős–Rényi network as the host contact network.** Results for six replicates are shown in separate plots. The solid black line represents the estimated effective population size over time, while the grey region indicates the credible intervals. The horizontal heatmap describes the coalescent event intensity; the darker the color, the greater the number of events occurring within a given time interval. Simulation details are in Section 5.6, supplementing Section 3.1.4.

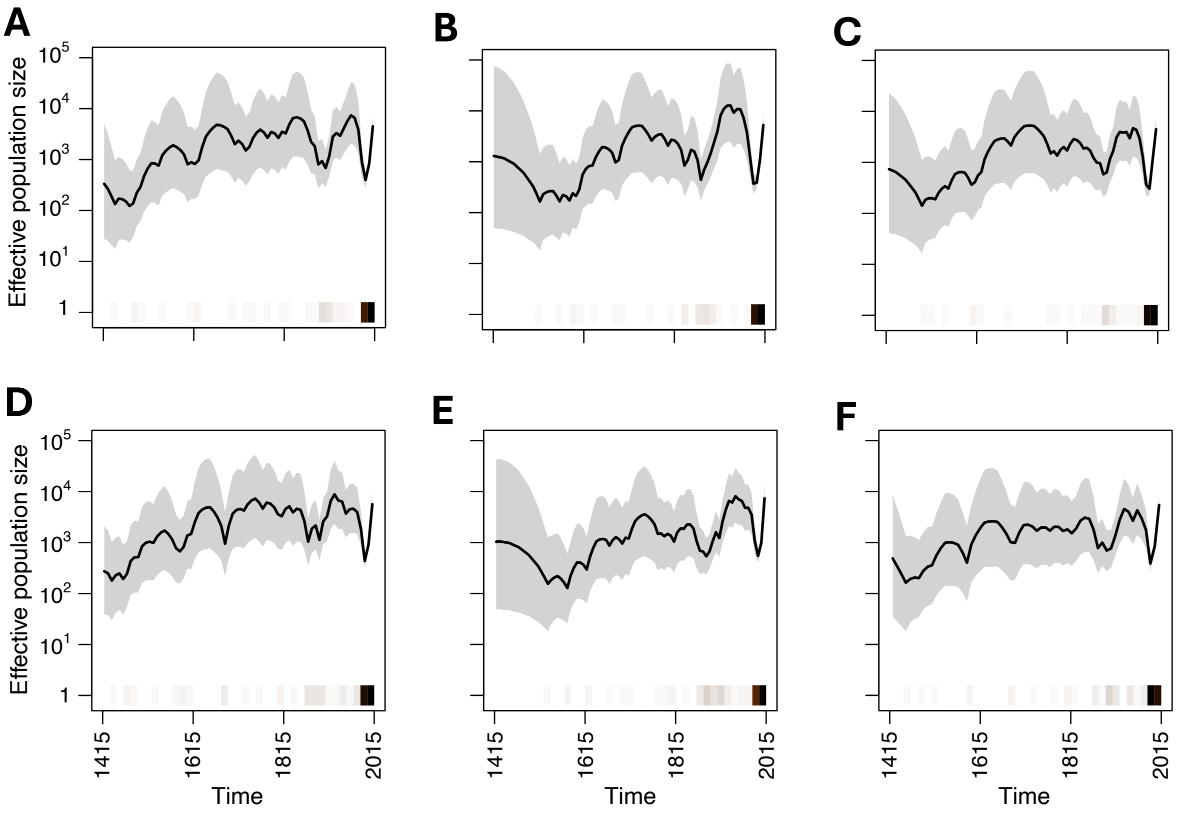


**Figure S7. Effective population size trajectory inferred by the BNPR method for the Karonga simulation using a Barabási–Albert network as host contact network.** Results for the six replicates are shown in separate plots. The figure design follows Figure S6. Simulation details are in Section 5.6, supplementing Section 3.1.4.

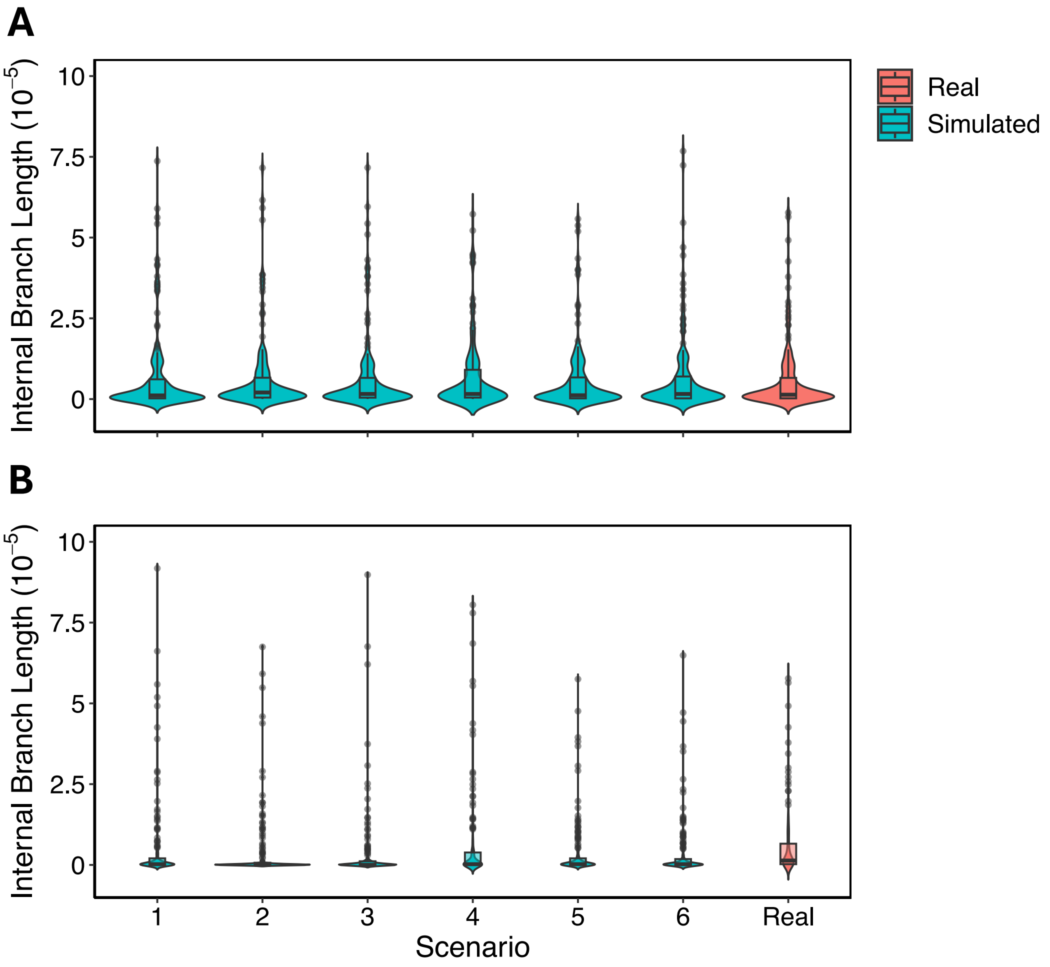


**Figure S8. Branch length distribution of genetic trees from sequences sampled in the last three years in simulated and Karonga datasets.** The green violins represent the branch length distributions of genetic trees reconstructed from samples for six replicates from simulations on different networks (Sections 3.1.4 and 5.6), while the red violins represent branch length distribution of genetic trees reconstructed from samples in the Karonga dataset. The x-axis numbers denote replicate IDs, and the “Real” represents the real dataset. One replicate from each contact network is shown in Figure 8 along with the real data. The violin for the real scenario is the same for A and B, with the violin width re-scaled for visualization purposes. (A) Simulated using the Erdős–Rényi network. (B) Simulated using the Barabási–Albert network.

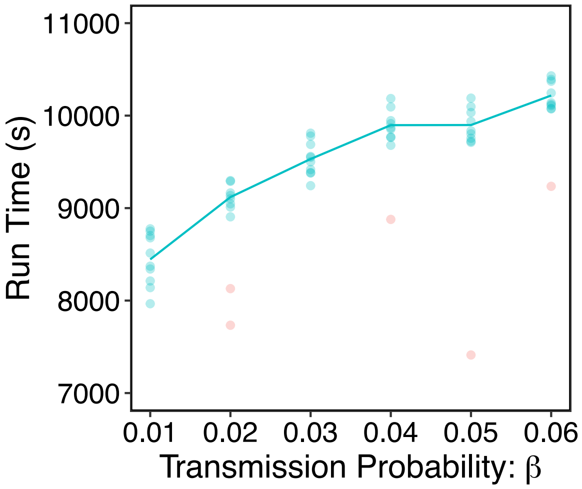


**Figure S9. Runtime benchmark for varying outbreak sizes by modifying base transmission probability ($\beta$) on Linux.** Each parameter set was run in ten replicates. Runtime for each replicate is represented by a dot, with the average runtime for each transmission probability shown by the line. Blue dots indicate successful replicates, where an outbreak occurred and persisted throughout the simulation. Red dots indicate unsuccessful replicates, where the outbreak died out before the simulation ended. The memory limit for these runs was set to 5GB.

**Configuration File S1. Configuration for running `OutbreakSimulator` for the SARS-CoV-2 simulation with two treatment stages.** Relevant files generated by the pre-simulation modules must be present in the base working directory `cwdir`. Descriptions of the parameters and simulation processes are provided in Section 3.1.1 and 5.3.

```
{
  "BasicRunConfiguration": {
    "cwdir": "covid",
    "n_replicates": 10
  },
  "EvolutionModel": {
    "subst_model_parameterization": "mut_rate",
    "n_generation": 730,
    "mut_rate": 1.8e-6,
    "within_host_reproduction": false,
    "within_host_reproduction_rate": 0,
    "cap_withinhost": 1
  },
  "SeedsConfiguration": {
    "seed_size": 1,
    "use_reference": true
  },
  "GenomeElement": {
    "use_genetic_model": true,
    "ref_path": "EPI_ISL_402124.fasta",
    "traits_num": {
      "transmissibility": 1,
      "drug_resistance": 2
    }
  },
  "NetworkModelParameters": {
    "use_network_model": true,
    "host_size": 10000
  },
  "EpidemiologyModel": {
    "model": "SEIR",
    "epoch_changing": {
      "n_epoch": 3,
      "epoch_changing_generation": [250, 500]
    },
    "genetic_architecture": {
      "transmissibility": [1, 1, 1],
      "cap_transmissibility": [10, 10, 10],
      "drug_resistance": [0, 1, 2],
      "cap_drugresist": [0, 10, 10]
    },
    transition_prob": {
      "S_IE_prob": [0.07, 0.07, 0.07],
      "I_R_prob": [0.035, 0.09, 0.08],
      "R_S_prob": [0.05, 0.15, 0.15],
      "latency_prob": [1, 1, 1],
      "E_I_prob": [0.3, 0.3, 0.3],
      "I_E_prob": [0, 0, 0],
      "E_R_prob": [0, 0, 0],
      "sample_prob": [0.0001, 0.0001, 0.0001],
      "recovery_prob_after_sampling": [0, 0, 0]
    },
```

```json
    "massive_sampling": {
      "event_num": 0,
      "generation": [],
      "sampling_prob": [],
      "recovery_prob_after_sampling": []
    },
    "super_infection": false
  },
  "Postprocessing_options": {
    "do_postprocess": true,
    "tree_plotting": {
      "branch_color_trait": 1,
      "heatmap": "drug_resistance"
      },
    "sequence_output": {
      "vcf": true,
      "fasta": false
      }
    }
  }
}
```

**Configuration File S2. Configuration for running `OutbreakSimulator` for *Mtb* simulations with various host contact network structures.** The network file specifying the contact structure and other relevant files generated by the pre-simulation modules must be present in the base working directory `cwdir`. Descriptions of the parameters and simulation processes are provided in Section 3.1.2 and 5.4.

```
{
  "BasicRunConfiguration": {
    "cwdir": "tb",
    "n_replicates": 10
  },
  "EvolutionModel": {
    "subst_model_parameterization": "mut_rate",
    "n_generation": 3650,
    "mut_rate": 3.12e-10,
    "within_host_reproduction": false,
    "within_host_reproduction_rate": 0,
    "cap_withinhost": 1
  },
  "SeedsConfiguration": {
    "seed_size": 5,
    "use_reference": false
  },
  "GenomeElement": {
    "use_genetic_model": true,
    "ref_path": "GCF_000195955.2_ASM19595v2_genomic.fna",
    "traits_num": {
      "transmissibility": 1,
      "drug_resistance": 0
    }
  },
  "NetworkModelParameters": {
    "use_network_model": true,
    "host_size": 10000
  },
  "EpidemiologyModel": {
    "model": "SEIR",
    "epoch_changing": {
      "n_epoch": 1,
      "epoch_changing_generation": []
    },
    "genetic_architecture": {
      "transmissibility": [1],
      "cap_transmissibility": [10],
      "drug_resistance": [0],
      "cap_drugresist": [1.5]
    },
    "transiton_prob": {
      "S_IE_prob": [0.003],
      "I_R_prob": [0.008],
      "R_S_prob": [0.05],
      "latency_prob": [1],
      "E_I_prob": [0.01],
      "I_E_prob": [0],
      "E_R_prob": [0],
      "sample_prob": [0.00002],
      "recovery_prob_after_sampling": [0]
    },
```

```
    "massive_sampling": {
      "event_num": 0,
      "generation": [],
      "sampling_prob": [],
      "recovery_prob_after_sampling": []
    },
    "super_infection": false
  },
  "Postprocessing_options": {
    "do_postprocess": true,
    "tree_plotting": {
      "branch_color_trait": 1,
      "heatmap": "drug_resistance"
      },
    "sequence_output": {
      "vcf": true,
      "fasta": false
      }
    }
  }
}
```

**Configuration File S3. Configuration for running `OutbreakSimulator` to simulate the SARS-CoV-2 outbreak in Haslemere.** Relevant files generated by the pre-simulation modules must be present in the base working directory `cwdir`. Descriptions of the parameters and simulation processes are provided in Sections 3.1.3 and 5.5.


```json
{
  "BasicRunConfiguration": {
    "cwdir": "Haslemere",
    "n_replicates": 10
  },
  "EvolutionModel": {
    "n_generation": 365,
    "mut_rate": 0,
    "subst_model_parameterization": "mut_rate_matrix",
    "mut_rate_matrix": [[0,7.165664751644342e-08,2.8379053325361 28e-07,5.446119130833203e-08],
        [1.1729557471338957e-07,0,3.5148079746603924e-08,1.5082154072511841e-06],
        [4.325598694008407e-07,3.27284006207541e-08,0,2.3283926854277938e-07],
        [5.050713646311029e-08,8.544838143470873e-07,1.4166857139695633e-07,0]],
    "within_host_reproduction": false,
    "within_host_reproduction_rate": 0,
    "cap_withinhost": 1
  },
  "SeedsConfiguration": {
    "seed_size": 16,
    "use_reference": false
  },
  "GenomeElement": {
    "use_genetic_model": true,
    "ref_path": "EPI_ISL_402124.fasta",
    "traits_num": {
      "transmissibility": 1,
      "drug_resistance": 0
    }
  },
  "NetworkModelParameters": {
    "use_network_model": true,
    "host_size": 12907
  },
  "EpidemiologyModel": {
    "model": "SEIR",
    "epoch_changing": {
      "n_epoch": 1,
      "epoch_changing_generation": []
    },
    "genetic_architecture": {
      "transmissibility": [1],
      "cap_transmissibility": [10],
      "drug_resistance": [0],
      "cap_drugresist": [0]
    },
    "transition_prob": {
      "S_IE_prob": [0.03],
      "I_R_prob": [0.015],
      "R_S_prob": [0.025],
      "latency_prob": [1],
      "E_I_prob": [0.15],
      "I_E_prob": [0],
```

```
      "E_R_prob": [0],
      "sample_prob": [0.0007],
      "recovery_prob_after_sampling": [0]
    },
    "massive_sampling": {
      "event_num": 0,
      "generation": [],
      "sampling_prob": [],
      "recovery_prob_after_sampling": []
    },
    "super_infection": false
  },
  "Postprocessing_options": {
    "do_postprocess": true,
    "tree_plotting": {
      "branch_color_trait": 1,
      "heatmap": "drug_resistance"
      },
    "sequence_output": {
      "vcf": true,
      "fasta": false
      }
    }
  }
}
```

**Configuration File S4. Configuration for running `OutbreakSimulator` to simulate the *Mtb* outbreak in Karonga.** The network file specifying the contact structure and other relevant files generated by the pre-simulation modules must be present in the base working directory `cwdir`. Descriptions of the parameters and simulation processes are provided in Section 3.1.4 and 5.6.


```json
{
  "BasicRunConfiguration": {
    "cwdir": "Karonga",
    "n_replicates": 6
  },
  "EvolutionModel": {
    "n_generation": 2433,
    "subst_model_parameterization": "mut_rate_matrix",
    "mut_rate_matrix": [[0,2.536844537141152e-10,9.179395141595392e-10,3.928840940389246e-11],
            [1.326691743031835e-10,0,1.772796094563627e-10,4.752540935024862e-10],
            [4.818131373558009e-10,1.7792918359788217e-10,0,1.31707708034985e-10],
            [3.9288409403892454e-11,9.08761026959088e-10,2.5092655253902036e-10,0]],
    "within_host_reproduction": false,
    "within_host_reproduction_rate": 0,
    "cap_withinhost": 1
  },
  "SeedsConfiguration": {
    "seed_size": 375,
    "use_reference": false
  },
  "GenomeElement": {
    "use_genetic_model": false,
    "ref_path": "GCF_000195955.2_ASM19595v2_genomic.fna",
    "traits_num": {
      "transmissibility": 1,
      "drug_resistance": 0
    }
  },
  "NetworkModelParameters": {
    "use_network_model": true,
    "host_size": 300000
  },
  "EpidemiologyModel": {
    "model": "SEIR",
    "epoch_changing": {
      "n_epoch": 2,
      "epoch_changing_generation": [2068]
    },
    "genetic_architecture": {
      "transmissibility": [0,0],
      "cap_transmissibility": [10,10],
      "drug_resistance": [0,0],
      "cap_drugresist": [1.5, 1.5]
    },
    "transition_prob": {
      "S_IE_prob": [0.004, 0.004],
      "I_R_prob": [0.008, 0.008],
      "R_S_prob": [0.01, 0.01],
      "latency_prob": [1, 1],
      "E_I_prob": [0.002, 0.002],
      "I_E_prob": [0, 0],
      "E_R_prob": [0, 0],
```

```
      "sample_prob": [0, 0.01],
      "recovery_prob_after_sampling": [0, 0]
    },
    "massive_sampling": {
      "event_num": 0,
      "generation": [],
      "sampling_prob": [],
      "recovery_prob_after_sampling": []
    },
    "super_infection": false
  },
  "Postprocessing_options": {
    "do_postprocess": true,
    "tree_plotting": {
      "branch_color_trait": 1,
      "heatmap": "drug_resistance"
      },
    "sequence_output": {
      "vcf": true,
      "fasta": false
      }
    }
  }
}
```
